# High-resolution two-photon transcranial imaging of brain using direct wavefront sensing

**DOI:** 10.1101/2020.09.12.294421

**Authors:** Congping Chen, Zhongya Qin, Sicong He, Shaojun Liu, Shun-Fat Lau, Wanjie Wu, Dan Zhu, Nancy Y. Ip, Jianan Y. Qu

**Author notes:** These authors contributed equally to this work (Congping Chen, Zhongya Qin). Corresponding author (J.Y.Q.).

## Abstract

Imaging of the brain in its native state at high resolution poses major challenges to visualization techniques. Two-photon microscopy integrated with the thinned-skull or optical clearing skull technique provides a minimally invasive tool for *in vivo* imaging of the cortex of mice without activating immune response and inducing brain injury. However, the imaging contrast and resolution are severely compromised by the optical heterogeneity of the skull, limiting the imaging depth to the superficial layer. Here, we develop adaptive optics two-photon microscopy for high-resolution transcranial imaging of layer 5 pyramidal neurons up to 700 μm below pia in living mice. In particular, an optimized configuration of imaging system and new wavefront sensing algorithm are proposed for accurate correction for the aberrations induced by the skull window and brain tissue. We investigated microglia-plaque interaction in living brain of Alzheimer’s disease and demonstrated high-precision laser dendrotomy and single-spine ablation.

## Introduction

Direct visualization and manipulation of neurons, glia and microvasculature in their native environment is crucial to understand how the brain functions. With the growth of fluorescent protein and transgenic technology, two-photon excited microscopy has become an indispensable tool for *in vivo* brain imaging of small rodents over recent decades because of its high spatial resolution and optical-sectioning capability(1, 2). However, the biggest obstacle for direct imaging of the brain in living animal is the opaque skull because it attenuates both the excitation and emission photons of two-photon microscopy severely, yielding poor image quality even in the superficial brain region.

The primary methods of providing optical access to the mouse brain are the open-skull and thinned-skull protocols(3, 4). The major limitation of open-skull approach is that skull removal will inevitably trigger the glia-mediated inflammatory reaction and disturb the neuronal physiology(5). Although an exceedingly thin skull could provide an imaging resolution close to that with an open-skull window at the superficial layer, the probability of mechanical disruption of cortex and activation of neuroinflammation is high and optical access is restricted to a very small area(4). Furthermore, an overly thinned skull is not suitable for chronic *in vivo* imaging because the newly grown skull must be constantly removed to ensure the optical quality of cranial window(4). To minimize the risk of brain trauma and inflammation, skull can be mechanically thinned to a certain thickness (~ 50 μm) to effectively reduce the scattering while holding its structural integrity to protect the underlying brain(6). Alternatively, optical clearing technique can improve the skull transparency by degrading the collagen fibers and removing the inorganic minerals with chemical reagents(7–9). However, aberrations arising in optical heterogeneity in the mechanically thinned or chemically cleared skull hamper transcranial brain imaging performance in both resolution and depth.

Adaptive optics (AO), originally developed for astronomical telescopes, has been introduced recently to improve two-photon microscopy by correcting system- or sample-induced aberrations(10, 11). The wavefront aberrations can be determined by either direct(12–16) or indirect(17–19) methods. The direct wavefront sensing approach employs a Shack-Hartman wavefront sensor (SHWS) to measure the wavefront distortion of the nonlinear fluorescence guide star. This method is fast, robust and photon-efficient, enabling two-photon imaging of layer 5 neurons through an open-skull cranial window(14, 15), and its application to the thinned skull window is limited to ~ 500 μm below the pia(15). In this work, we developed AO two-photon microscopy for high-resolution cortical imaging through both thinned-skull and optical clearing skull windows (**Fig. S1**). We built an ultra-sensitive SHWS incorporating a microlens array and an electron-multiplying charge-coupled device (EMCCD) to measure the wavefront of a descanned two-photon excited fluorescent (TPEF) guide star. The wavefront distortion was fed to a deformable mirror (DM) to introduce a compensating distortion to the excitation light, correcting the aberrations. We optimized the excitation numerical aperture (NA) of the microscope system, which alleviated the scattering of the excitation laser and also extended the depth of direct wavefront sensing. We advanced the wavefront sensing algorithm by averaging the Shack-Hartman images from arbitrarily distributed near-infrared (NIR) guide stars in a three-dimensional (3D) subvolume, allowing the reliable determination of aberration beneath the skull window and brain tissue. Using this system, we first characterized the optical properties of the skull windows and then achieved *in vivo* neuronal imaging in mouse brains with much improved resolution and signal intensity up to ~700 μm below pia. We then investigated the interaction between microglia and plaque in a mouse model of Alzheimer’s disease (AD). Taking advantage of the tight focus provided by AO correction, we demonstrated precise laser-mediated dendrotomy and single-spine ablation of layer 5 pyramidal neurons, and studied the microglial dynamic response to this neuronal microsurgery.

## Results

### Optimization of the imaging system configuration and new wavefront sensing algorithm

We first optimized the microscope system for transcranial deep brain imaging using a reduced NA excitation and a high NA collection configuration. This approach aimed to mitigate optical scattering and aberrations of the excitation laser from the skull window and brain tissue while maintaining a high collection efficiency of fluorescence signal for imaging. As shown in **Fig. 1a**, the excitation NA could be reduced by underfilling the back aperture of a high NA objective. To evaluate the effectiveness of this approach, we conducted *in vivo* imaging of YFP-labelled neurons (Thy1-YFP mice) through a thinned-skull window of 50 μm thickness with gradually decreasing the excitation NA from 1.05 to 0.7. Despite the decrease of effective NA in the underfilled configuration, we did not observe obvious degradation of the imaging resolution even in the topmost cortical layers (**Fig. S2**). This should be attributed to the high-NA rays being highly scattered by the skull/brain tissue and contributing little to the focus. Under the same excitation power, the fluorescence intensity was enhanced up to twofold when reducing the excitation NA from 1.05 to 0.7 (**Fig. 1b-c**). However, with further reduction of excitation NA, the imaging resolution decreased (> 0.6 μm) and was not sufficient to visualize the fine structures such as dendritic spines. These results indicated that using an underfilled objective is beneficial for deep-brain imaging through a skull window, in agreement with another study of the open-skull preparation(20). More importantly, because it is only necessary to correct aberrations in the excitation path for two-photon microscopy, the reduction of focal cone angle can also alleviate the scattering of the guide star signal, improve the quality of the Shack–Hartmann spot image and enabled wavefront measurement at deeper region (**Fig. 1d-e** and **Fig. S3**). The benefits are attributed to that SHWS favorably collects the descanned TPEF guide star signals from the reduced focal cone angle of excitation beam and effectively rejects scattered fluorescence from higher NA. To further extend the depth of wavefront measurement, we investigated how a NIR guide star could improve direct wavefront sensing through a thinned-skull window(15). We labelled the microvasculature with Evans Blue by using retro-orbital injection into Thy1-YFP mice. Evans Blue can be excited efficiently with a 920 nm laser and emits fluorescence at 680 nm. We compared the guide star images in SHWS generated using both kinds of fluorophores at the same location. As shown in **Fig. S4**, Evans Blue provides much better guide star images than YFP at all imaging depths, because of the reduced scattering at the longer wavelength fluorescence emission.

**Figure 1.**
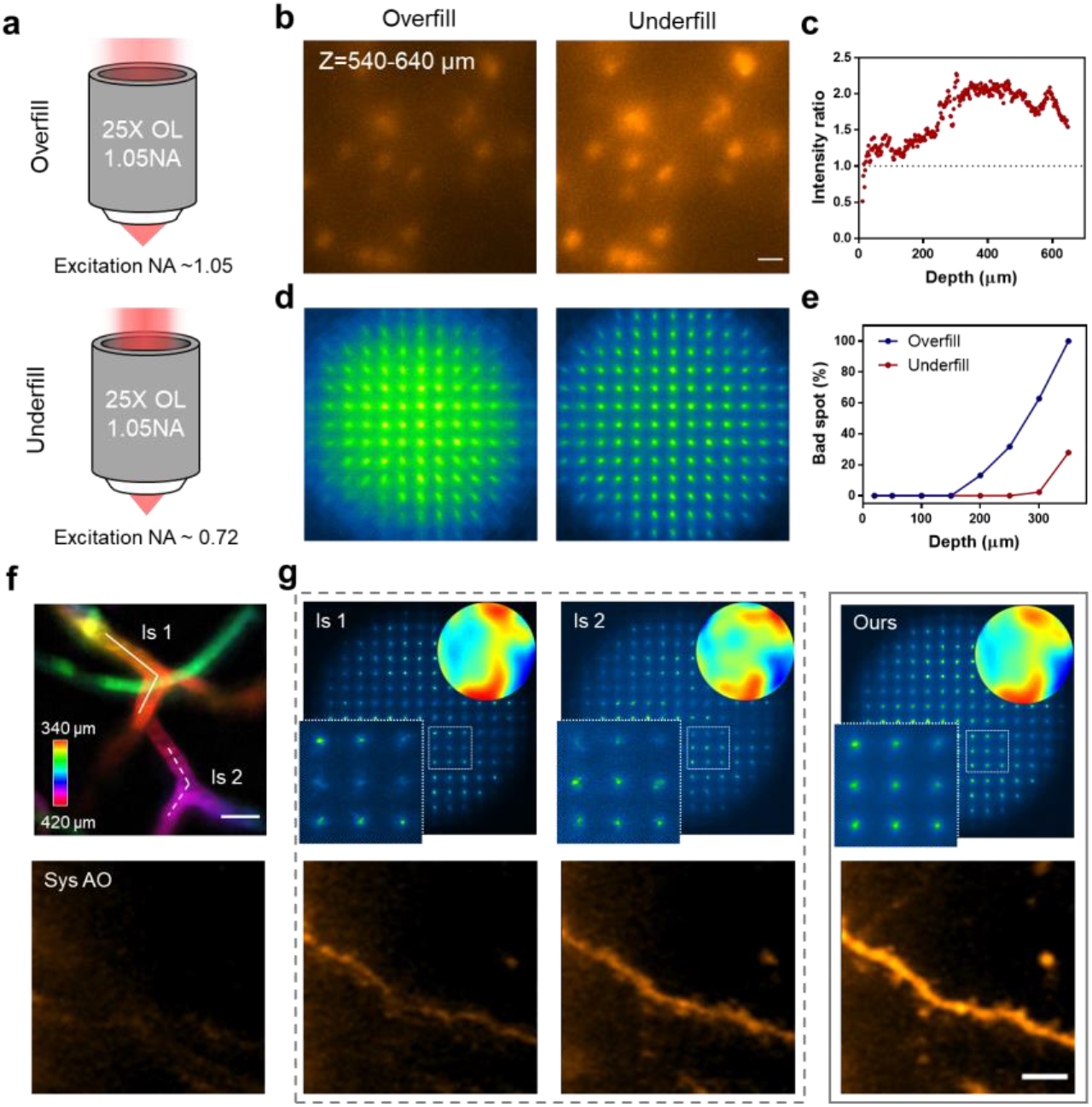
Optimization of the excitation NA and new wavefront sensing algorithm for transcranial brain imaging. **(a)** Schematic illustration of a high-NA objective in the overfilled (top) and underfilled (bottom) configuration. **(b)** xy maximum-intensity projection (MIP) images of the pyramidal neurons in Thy1-YFP mice through a 50-μm thinned-skull window acquired using the overfilled (left) and underfilled (right) objective. Imaging depth range: 540-640 μm. Scale bar: 20 μm. Details were presented in Fig. S2. **(c)** Enhancement of signal intensity with depth by reducing the excitation NA. The intensity ratio is the average intensity of the brightest 0.3% pixels in the xy image acquired with the underfilled objective divided by that with the overfilled objective as shown in Fig. S2a. **(d)** Representative guide star images of YFP fluorescence in Thy1-YFP mice when the objective was overfilled (left) and underfilled (right) at 250 μm below the thinned skull. Details are shown in Fig. S3. **(e)** Percentages of bad spots in the Shack-Hartman spot image for overfilled (blue) and underfilled (red) configuration. A bad spot is the one with poor signal quality to make its center unidentifiable. **(f)** Top: Depth-coded images of the NIR-dye labeled microvascular vessels for direct wavefront sensing. Two segments of vessels at different depths (solid line and dashed line labelled with ls 1 and ls 2) were line scanned for wavefront measurement. Scale bar: 10 μm. Bottom: two-photon images of YFP labelled dendrite with system correction only. **(g)** Top row: guide star images on the SHWS with only ls 1 (left) or ls 2 (middle) and our algorithm (right). The left-bottom corners show magnified views of the boxed regions and the right-top corners display the corrective wavefront pattern on the DM. Bottom row: the corresponding AO corrected images. Details were presented in Fig. S7. Scale bar: 5 μm.

Since the skull is highly heterogeneous and contributes for the major aberrations, we characterized the optical aberrations induced by the thinned-skull window in an *in vitro* preparation. We created a 3D tissue phantom by dispersing fluorescent beads (0.2 μm in diameter) in a mixture of Evans blue/agarose and then placing a piece of isolated thinned-skull (50 μm in thickness) on top of the sample. Evans Blue fluorescence provided bright and uniform guide stars for direct measurement of the aberrations of the thinned-skull window, while the fluorescent beads were used to evaluate the PSF distortion caused by the aberrations. Because the wavefront distortion varies spatially due to the skull heterogeneity, we first investigated the isoplanatic FOV within which the aberrations were similar. We performed AO corrections by averaging the aberrations over a series of FOV ranges and compared the enhancement of the fluorescence intensity of the central bead (**Fig. S5**). As can be seen, the aberrations would average out when the guide star was scanned over a too large field, while if the scanned field is too small, tissue scattering yields irregular Shack–Hartmann spots and induces errors in determining the aberration(13). The optimal scanning FOV was found to be a square with sides of 30~60 μm (**Fig. S5b**). Further, we characterized the aberrations of the thinned-skull window at various depths (**Fig. S6**). The results showed that AO increased the fluorescence intensity up to 10-fold and restored near-diffraction-limited resolution over 600 μm below the skull.

Next, we applied the AO approach to *in vivo* imaging of the mouse cortex in Thy1-YFP mice. Evans blue was retro-orbitally injected to label the brain vasculature and served as the guide star in the deep brain region. To measure the aberrations, we excited the labelled microvessels in 60×60 μm^2^ and integrated the de-scanned fluorescence signal on the SHWS. However, because the brain tissue and the overlying skull were so heterogeneous, the Shack–Hartmann spots became irregular and asymmetrical even when the guide star signal was scanned over a segment of blood vessel (**Fig. 1f-g**). This induced large errors in spot center identification and wavefront measurement, resulting in inaccurate or incomplete AO correction. Given that an isoplanatic correction is valid within a small 3D volume (60×60×60 μm^3^), not merely in a 2D focal plane, we developed a wavefront reconstruction algorithm by summing the SHWS images captured at different depths within the isoplanatic volume and then spatially filtering each spot with its neighbors (see method). This approach yields a clear image of Shack–Hartmann spots and single spots stand out in each cell of the SHWS, enabling more accurate aberration determination (**Fig. 1g**). By using this algorithm, we corrected the aberrations reliably and improved the imaging performance (**Fig. 1g** and **Fig. S7**).

### High-resolution cortical imaging through the thinned-skull window

Taking the advantage of the optimized imaging system, NIR guide star and new algorithm of wavefront sensing, we conducted *in vivo* transcranial imaging of cortex through a thinned skull of ~ 50-μm thickness in Thy1-YFP mice. As can be seen, even with system aberration corrected, the neuronal dendrites and somata were severely blurred by aberrations caused by skull and brain tissue (**Fig. 2a-c**). With full AO correction, however, pyramidal neurons spanning hundreds of microns in depth could be resolved clearly, along with the surrounding microvessels. Quantitative comparisons show that AO not only dramatically enhanced the fluorescence intensity, but also recovered the optimal imaging resolution at depths as great as 680 μm below the pia (**Fig2. a-c** and **Fig. S8**). These results lead to the conclusion that AO is essential and efficient for high-resolution and deep-brain imaging through minimally invasive thinned-skull windows.

**Figure 2.**
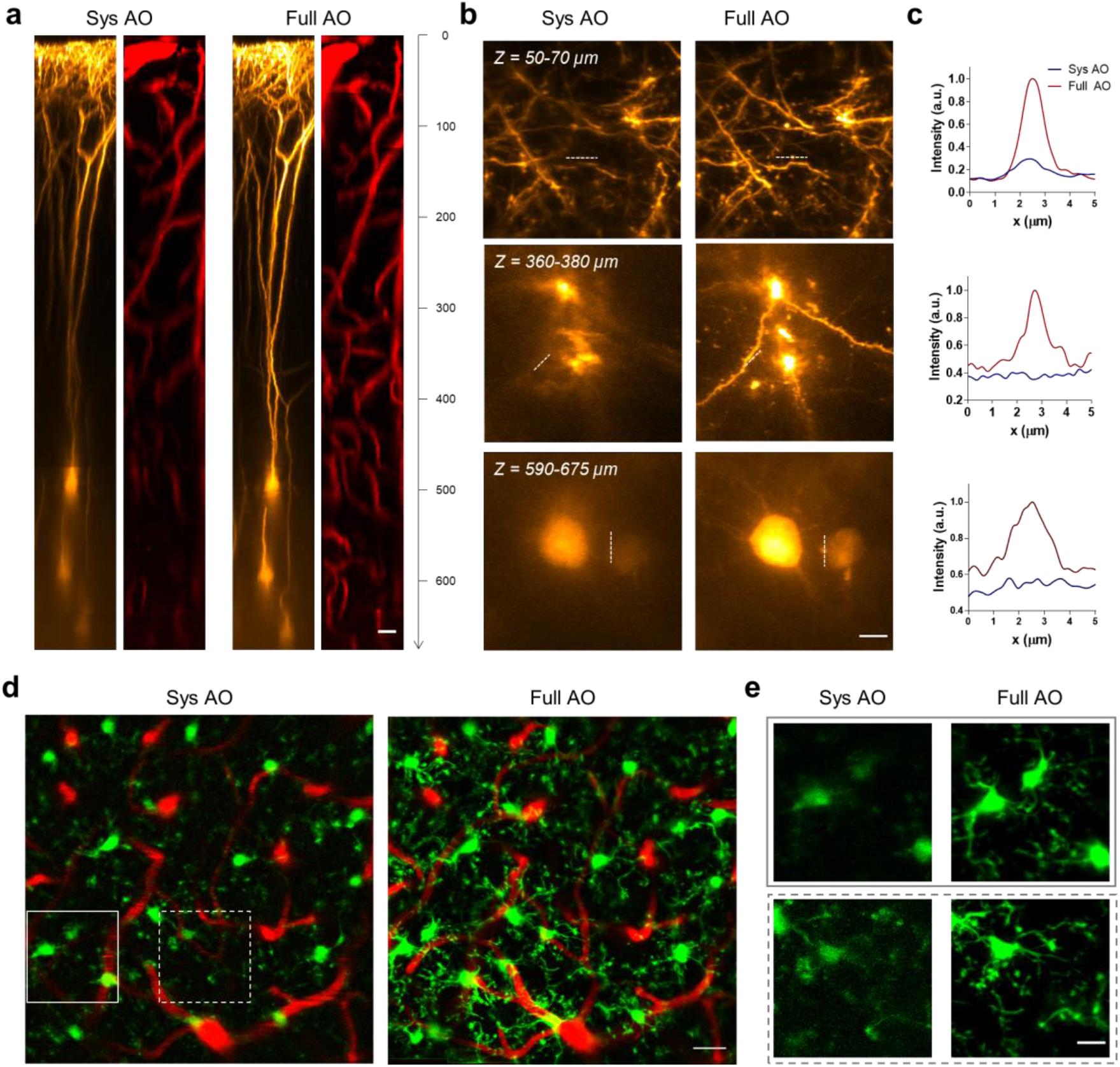
*In vivo* AO imaging of the brain at high resolution through a thinned-skull window. **(a)** xz MIP images of the pyramidal neurons (orange) and microvasculature (red) in Thy1-YFP mice through a thinned-skull window (50 μm in thickness) with system correction only (left) and full AO correction (right). AO correction was performed every 50 μm of depth. Scale bar: 20 μm. **(b)** xy MIP of the stack images in (a). Scale bar: 5 μm. **(c)** Intensity profiles along the dashed lines in (b) with system (blue) and full (red) AO correction. **(d)** *In vivo* imaging of microglia (green) and microvessels (red) at 350-400 μm below the pia in the Cx3Cr1-GFP mice with system (left) and full (right) AO corrections. Full AO correction were performed every 40 μm and 5×5 subregions were stitched together to form the entire image. Scale bar: 20 μm. **(e)** Magnified views of the boxed region in (d). Scale bar: 10 μm.

Microglia, the brain-resident phagocytes, play a critical role in brain homeostasis and neurological diseases. The resting microglia with motile processes are highly sensitive to subtle changes in brain parenchyma, and can become activated rapidly with substantial changes in morphology and function upon brain damage or injury(21). Therefore, minimally invasive imaging tools with the ability to resolve the fine processes are crucial for the study of microglial physiology in the native environment. Taking advantage of our approach, we conducted *in vivo* imaging of Cx3Cr1-GFP mice with microglia labeled with green fluorescence protein (GFP). We first examined whether the thinned-skull (50-μm thickness) preparation triggered the inflammatory response of microglia using time-lapse imaging. Here, ramification and surveillance of the microglial processes were quantified and monitored following the thinned-skull surgery to evaluate the potential surgical effect on microglia activation(22). As indicated by the ramified morphology and surveying behavior (**Fig. S9**), the microglia were not activated and the “thick” thinned skull window effectively protected underlying brain tissue. By virtue of the fast AO correction, we sequentially measured and corrected the aberrations in each subvolume and then stitched them together to form a mosaic image of large FOV. As can be seen, after full AO correction, the branching processes of microglia can be visualized clearly across the entire FOV 400 μm below the pia (**Fig. 2d-e**).

### High-resolution cortical imaging through the optical clearing skull window

The optical clearing skull window is another technique for minimally invasive imaging of the brain(7–9). By degrading the collagens and inorganic minerals with chemical reagents, the scattering of the mouse skull can be reduced greatly, enabling *in vivo* imaging of the underlying cortex without disturbing brain homeostasis (**Fig. 3a** and **Fig. S10–11**). However, although the fluorescence intensity was enhanced tremendously after optical clearing, the imaging contrast and resolution were still low because of the skull-induced aberration (**Fig. 3a**). Following the study of AO imaging through thinned skull window, we sought to explore whether our AO approach could also improve imaging performance through optical clearing windows. The *in vitro* imaging results showed that despite the large aberration caused by the heterogeneity of skull and refractive index mismatch between the clearing reagents and water, our AO approach can recover the optimal imaging resolution effectively up to 500 μm below the skull (**Fig. S12**). *In vivo* imaging of the mouse cortex shows that AO improved the imaging resolution and fluorescence brightness up to 600 μm below the pia (**Fig. 3b-c**). Further, the branching processes of microglia can also be visualized clearly with AO correction (**Fig. 3d-e**), which allows us to study the dynamics of microglial processes in both physiological and pathological conditions. It should be noted that the imaging depth of the optical clearing skull window is smaller than the thinned skull preparation, likely due to the larger skull-induced scattering.

**Figure 3.**
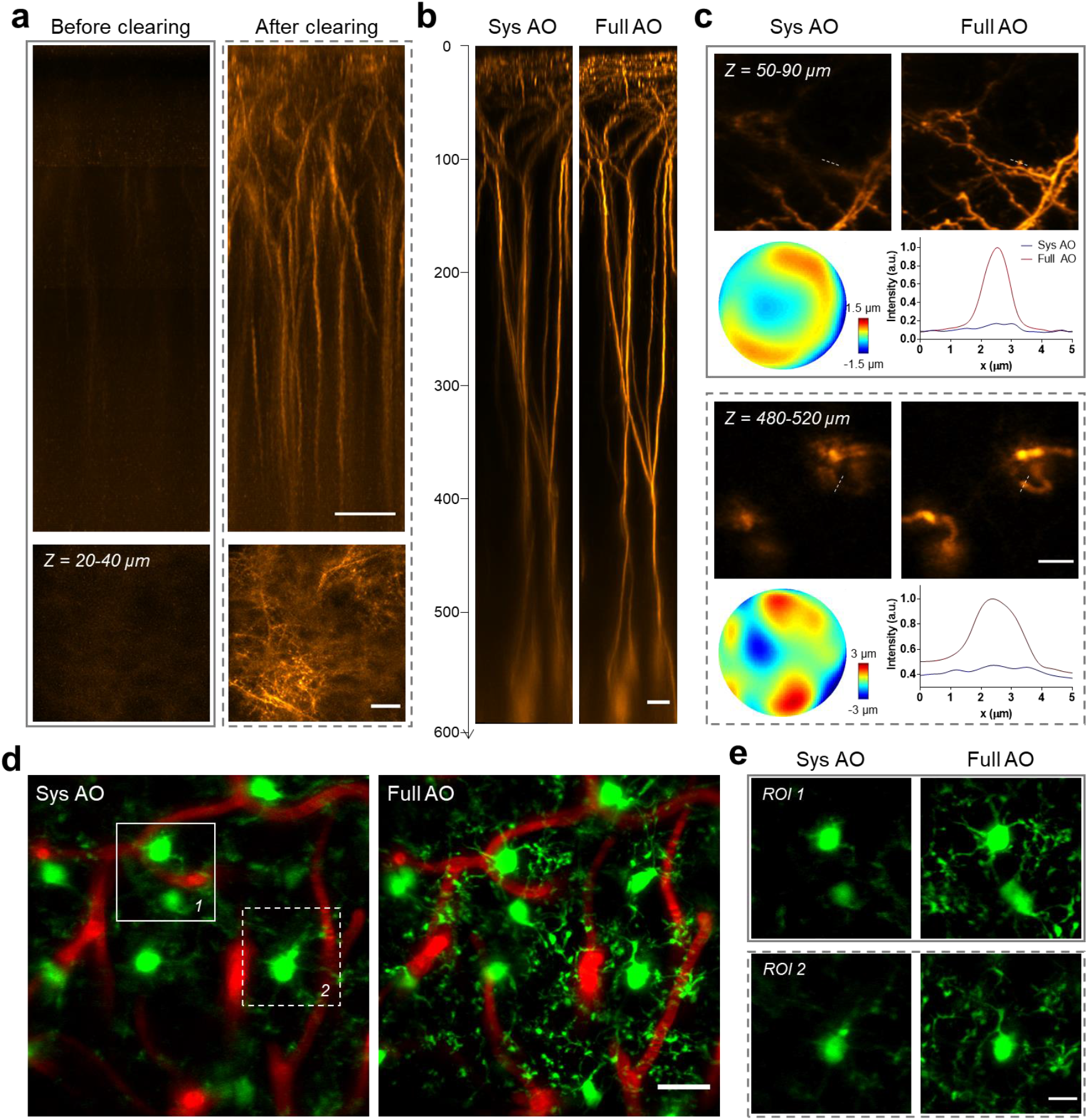
AO recovers high-resolution imaging of the cortex through an optical clearing window. **(a)** xz MIP of two-photon images of the YFP labelled neurons in the Thy1-YFP mice before (left) and after (right) optical clearing. Scale bar: 50 μm. Bottom row shows the xy MIP of the stack images. Scale bar: 20 μm. **(b)** xz MIP images of the pyramidal neurons in Thy1-YFP mice through the optical clearing window with system (left) and full (right) AO correction. AO correction was performed at every 50 μm depth. Scale bar: 20 μm. **(c)** xy MIP of the stack images in (b) at two representative depths and the corresponding corrective wavefront and intensity profiles along the dashed lines. Scale bar: 10 μm. **(d)** *In vivo* imaging of microglia (green) and microvessels (red) at 175-225 μm below the pia in the Cx3Cr1-GFP mice with system (left) and full (right) AO corrections. Full AO correction were performed every 40 μm and 3×3 subregions were stitched together to form the entire image. Scale bar: 20 μm. **(e)** Magnified views of images of the boxed regions in (d). Scale bar: 10 μm.

### High-resolution imaging of microglia-plaque interaction and high-precision laser surgery

By using our AO two-photon microscope, we can perform high-resolution transcranial imaging in deep cortical layers without interrupting the brain homeostasis, which is crucial for the study of microglial roles with laminar characteristics under certain pathological conditions. For example, we investigated the microglial activity in the AD mice brain which has a laminar distribution of amyloid plaques(23). We observed significant morphological and functional differences between the plaque-associated and plaque-distant microglia in layer II/III of mice cortex (**Fig. 4a-b**). While the plaque-distant microglia has highly ramified processes with similar motility as that in the normal brain, the microglia surrounding the amyloid plaques shows less ramified morphologies with no obvious dynamics (**Fig. 4b**). These results indicate that normal microglia may undergo phenotype alteration that is associated with the layer-specific distribution of amyloid plaque during the progression of AD pathology. Our AO-assisted minimally invasive imaging method can also facilitate the therapeutic study on the microglia-mediated inflammation in AD.

**Figure 4.**
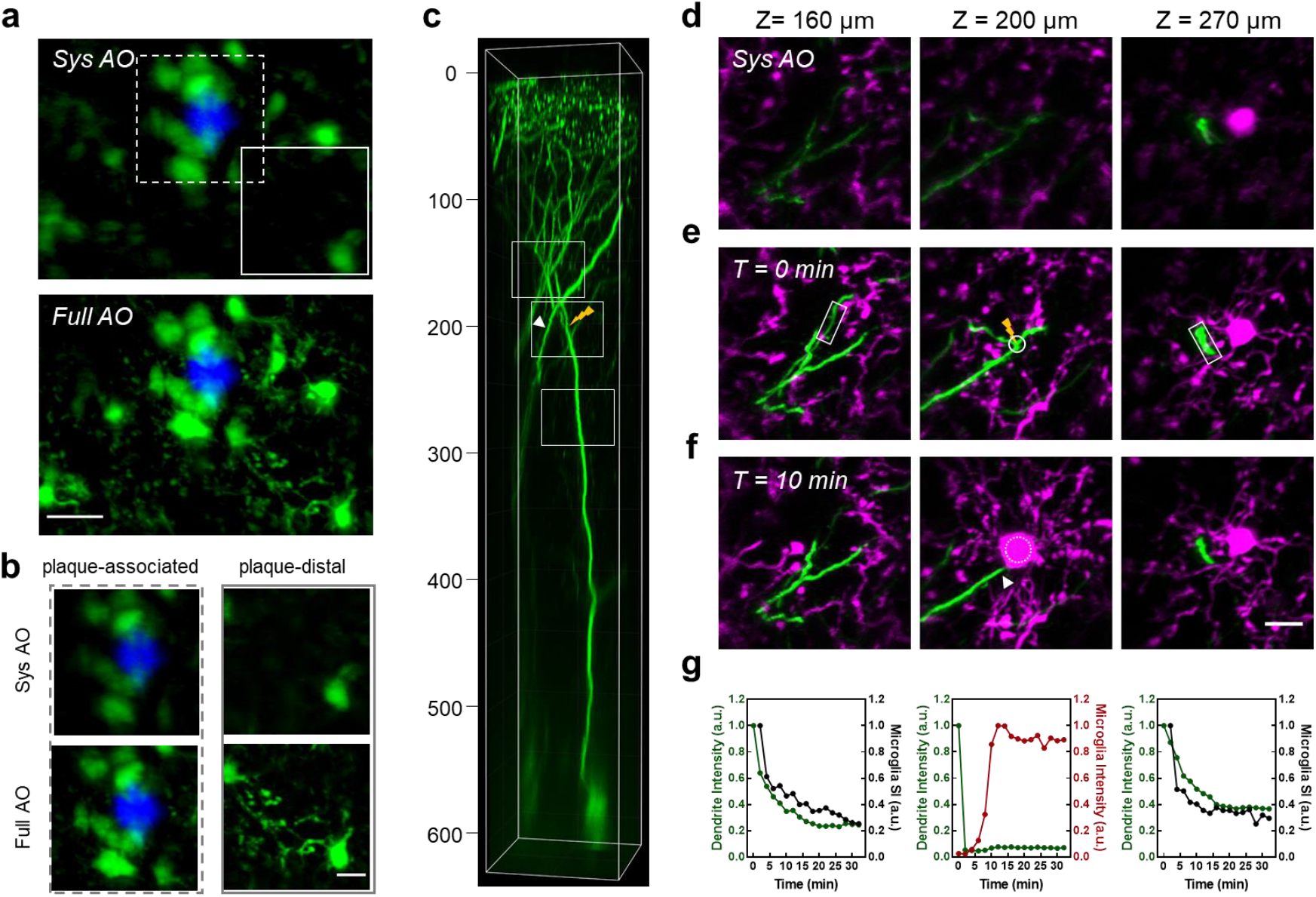
Study of microglial activities in AD mice and neuron-microglial interactions following precise laser micro-lesion. **(a)** *In vivo* imaging of amyloid plaque(blue) and microglia (green) at 230-270 μm below the pia in the APP-PS1/Cx3Cr1-GFP mice through a thinned-skull window with system (top) and full (bottom) AO corrections. Full AO correction was performed every 40 μm and 3×3 subregions were stitched together to form the entire image. Amyloid plaque was labeled by MeO-X04 through intraperitoneal injection. Scale bar: 20 μm. **(b)** Magnified views of the boxed region in (a) showing the plaque-associated (left) and plaque-distal (right) microglia. Scale bar: 10 μm. **(c)** 3D reconstruction of pyramidal neurons in Thy1-YFP/Cx3Cr1-GFP mice through a thinned-skull window with full AO correction. **(d-e)** xy MIP images of neuronal dendrites (green) and microglia (magenta) of the boxed region in (c) with system (d) and full (e) AO correction before laser ablation. Imaging depths: left column, Z = 140-180 μm; middle column, Z = 180-220 μm; right column, Z = 250-290 μm. The bifurcation point of the apical dendrite (Z = 200 μm) was targeted for laser microsurgery (thunder symbol in (c) and middle panel of (e)). **(f)** Full AO corrected images taken 10 minutes after laser ablation. The persistence of a nearby dendrite (arrowhead in (c) and the middle plane of (f)) indicates the confinement of the laser injury. Scale bar: 10 μm. **(g)** Characterization of microglial response to the dendritic degeneration. Left and right: microglial surveillance index (black line) and fluorescence decay (green line) of the distal and proximal dendrites (boxed region in the left and right panel of (e)); middle: influx of microglial processes to the injured spots represented by the microglial fluorescence intensity (dotted circle in the middle panel of (f)) and the fluorescence decay of the injured dendrite (solid circle in the middle panel of (e)).

Laser microsurgery, because of its high spatial precision of injury, has become a valuable tool for studying the cellular mechanisms that underlie various pathological phenomena such as neuronal degeneration and vascular disruption(24–27). However, high-precision laser surgery through skull windows is challenging because of large distortion of laser focus. We then applied the AO approach to study microglia-neuron interactions following laser-mediated neuronal injury through a thinned-skull window in Cx3Cr1-GFP/Thy1-YFP mice. We specifically targeted the first bifurcation point of the primary apical dendrite of a layer 5 pyramidal neuron and its neighboring compartments (**Fig. 4c**). As shown in **Fig. 4d-e**, the microglial processes and neuronal dendrites/ spines were clearly visualized using full AO correction. More importantly, AO also enabled precise laser microsurgery through the thinned-skull window, allowing us to ablate the branch point without influencing the nearby neurites, which is impossible otherwise. Time-lapse imaging revealed that the activation of microglia was highly correlated with the degeneration of the injured neuron (**Fig. 4f-g** and **Fig. S13**). While the local microglia near the injured site extended their processes rapidly towards the ablation point and completely wrapped around it within 10 minutes, microglia further away (either upper or deeper) also showed coordinated responses to the neuronal degeneration as indicated by decreased process motility (**Fig. 4f-g, Fig. S13** and **Movie. S1**). In addition, we also performed laser dendrotomy on a tuft dendrite of the layer 5 pyramidal neuron and high-precision single spine ablation without damaging the dendritic shaft and nearby spines, which resulted in distinct microglial responses (**Fig. S14** and **Movie. S2**). Although there is a chance that nearby unlabeled brain structures were injured, the removal of a single spine without influencing the parent dendrite indicates the submicron precision of AO-assisted laser microsurgery. These results demonstrated the great potential of AO for precise optical manipulation, in addition to high-resolution imaging.

## Discussion

In this work, we advanced the imaging system and AO technique for high-resolution deep-brain imaging through minimally invasive skull windows by employing a NIR guide star within the microvasculature. In particular, we optimized the excitation NA of our microscope system and showed that the use of an underfilled objective can not only improve the excitation efficiency, but also benefits direct wavefront sensing of TPEF guide stars. By scanning the NIR guide star within the microvasculature over a 3D subvolume and subsequently employing a wavefront processing algorithm, we achieved *in vivo* morphological imaging of layer 5 pyramidal neurons up to 700 μm below the 50-μm-thickness skull. Further, by taking advantage of the optimal point spread function provided by AO correction, we performed precise laser ablation of a single dendrite or spine and studied the interaction between neurites and microglial cells following neuronal microsurgery. Our results demonstrate that AO promises to advance the minimally invasive imaging tools and facilitate neuroscience research in the living brain.

Because of the extreme optical inhomogeneity of skull bone, the aberration varies quickly at different focal positions and the optimal corrective FOV is 30~60 μm for the thinned-skull and optical clearing skull windows, which is smaller than that of the open-skull cranial window (100 ~ 150 μm(14, 15, 28)). Thanks to fast AO correction by direct wavefront sensing, we measured the aberrations sequentially for each subvolume and stitched these subimages together to form an image of an entire large FOV. We have demonstrated that by using a NIR guide star with an emission peak at ~ 680 nm, direct wavefront sensing enabled imaging up to 700 μm below the pia through a thinned skull window. The imaging resolution and contrast in deeper brain region are still compromised by the dominant scattering caused by the skull and brain tissue. In this case, longer excitation/emission wavelengths and even three-photon absorption process are prefered(29, 30). In conjunction with emerging NIR fluorescent agents such as quantum dots and organic conjugated polymer dots(31, 32), we expect that direct-wavefront-sensing based AO can further increase the depth limit of minimally invasive skull windows.

## Materials and Methods

### Adaptive optics two-photon microscopy

A schematic diagram of our microscopy system is shown in **Fig. S1**. The 2P excitation beam (920 nm) from a tunable mode-locked femtosecond laser (Chameleon Ultra II, Coherent) was expanded and collimated by a pair of achromatic lens to slightly overfill the aperture of the DM (DM97-15, Alpao). The reflected beam with shaped wavefront was then compressed by a 4*f* telescope formed by two VIS-NIR achromatic doublets L5 and L6 (49-365 and 49-794, Edmunds) to match the aperture of a Galvo X scan mirror, which was conjugated with the DM. The Galvo XY-scan mirrors (6215H, Cambridge Technology) were mutually conjugated through a 4*f* relay formed by L7 and L8, both of which consist of two doublets (49-392, Edmunds). The Galvo Y and the rear pupil of the water-immersive objective (XLPLN25XSVMP2, ×25, 1.05 NA and 2mm working distance, Olympus) were then conjugated by the scan lens L9 and the tube lens L10 operating in the 4*f* relay configuration. Two groups of scan/tube lens combinations with magnification 3.33-fold or 2.25-fold were chosen to overfill or underfill the objective. For the overfilled condition, L9 consists of two doublets (49-391, Edmunds) and L10 is doublets (49-393, Edmund); for the underfilled condition, L9 consists of two doublets (49-392, Edmunds) and L10 is changed to doublets (49-365, Edmund). The objective was mounted on a motorized linear actuator (LNR50SEK1, Thorlabs) for axial sectioning. For specific imaging conditions requiring two excitation wavelengths simultaneously, another excitation beam (800 nm) from a tunable mode-locked femtosecond laser (Mira 900, Coherent) was integrated into the microscope system via a polarizing beam splitter. The system can operate in two modes: two-photon imaging and wavefront sensing.

For two photon imaging, the epi-fluorescence collected by the objective was reflected by a dichroic beam splitter D2 (FF757-Di01-25×36, Semrock) and directed to the photon detection unit. An interchangeable dichroic beam splitter D3 (FF560-Di01-25×36 or FF518-Di01-25X36, Semrock) was inserted at 45° to the beam path to separate the fluorescence into two current photomultiplier (PMT) modules (H11461-03 and H11461-01, Hamamatsu). Two band-pass filter F2 (HQ675/50M, Chroma or FF01-525/50, Semrock) and F3 (FF01-525/50 or FF03-447/60, Semrock) were placed before the PMTs to select the wavelength bands of the fluorescence. The PMTs current were then converted to voltage by two transimpedance amplifiers (SR570, Stanford Research and DLPCA-200, Femto) and subsequently fed into a multifunction data acquisition device (PCIe-6353, National Instrument). Custom-written C# software running in Visual Studio (Microsoft) was used to control the scanner/actuator and to acquire the TPEF images.

For wavefront sensing, the fluorescence emission from the guide star is transmitted through another dichroic beam splitter (Di02-R488-25×36, Semrock) replacing the D2 and then descanned by Galvo XY mirrors. The fluorescence signal was then reflected by the DM and separated from the excitation laser by the dichroic beam splitter D1 (FF705-Di01-25×36, Semrock), before being relayed by a lens pair L9 and L10(AC254-200-A and AC254-100-A, Thorlabs) to the microlens array (18-00197, SUSS MicroOptics) of the SHWS. The SHWS camera (iXon Ultra 888, Andor) was placed at the focal plane of the microlens array to record the pattern of spots of the fluorescent guide star, enabling direct measurement of its wavefront distortion. A bandpass filter F1 (HQ675/50M, Chroma or FF01-525/50, Semrock) was put before the SHWS to select the wavelength of the guide star signal. It should be noted that the DM, Galvo X and Y mirrors, rear pupil of the objective and microlens array of SHWS were all mutually conjugated. The DM and SHWS were operated in a close-loop configuration and controlled by a custom Matlab program integrated with the C# imaging software.

### Calibration of the DM

DM calibration following the previous procedure described by Wang was conducted before it was integrated into the imaging system(14). Briefly, the influence function of each DM actuator was measured using a Michaelson interferometer. The actuators’ driving voltages for the first 65 Noll’s Zernike modes were then obtained using the measured influence functions. After calibration, the DM can take any desired shape using a linear combination of these Zernike modes.

### System AO correction

Before any imaging experiment, the aberrations induced by the optical imperfections in the microscope system were corrected based on a sensorless AO algorithm(28). In brief, the TPEF intensity of a fluorescent dye (Rhodamine 6G) was used as a feedback signal to optimize the DM shape pattern. Seven to nine different values for each Zernike mode were applied sequentially to the DM, and the corresponding intensity of fluorescence was fitted to a Gaussian function to determine the optimum value for each Zernike mode. The first 21 Zernike modes (tip, tilt and defocus excluded) were used in the optimization cycle to determine and compensate for the system aberration ***Z***_sys_.

### Calibration of the SHWS

The SHWS was calibrated with the DM in the microscope system as described previously(16). Briefly, the nonlinear guide star from the fluorescent dye solution was descanned and used for AO calibration of the DM and SHWS in a close-loop configuration. The first 65 Zernike modes with root-mean-square *c_i_* and −*c_i_* were applied sequentially to the DM and the corresponding spot patterns 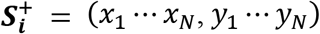 and 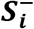 on the SHWS were recorded, where (*x_j_*, *y_j_*) represents the center location of the *j*-th spots. Then the influence matrix ***M***_sz_ of the DM to SHWS can be acquired, where

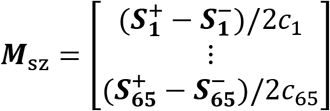

Each row of ***M***_sz_ represent the shift (x and y) of the spots on the SHWS to each Zernike mode. The 65 rows in the influence matrix ***M***_sz_ form the calibration basis for subsequent AO correction.

### Full AO correction

Full AO correction compensates both system- and sample-induced aberrations. First, the DM was set to correct the system aberration ***Z***_sys_ obtained using the sensorless method mentioned above. The TPEF signal of rhodamine at the FOV center creates a reference spot pattern on the SHWS ***S***_Sref_ = (*x*_1_ ⋯ *x_N_*, *y*_1_ ⋯ *y_N_*). To measure and correct the sample-induced aberration, a small FOV within the sample was scanned by the excitation laser to create a descanned and integrated wavefront on the SHWS. The SHWS images were first cross-correlated with a Gaussian function that equals to the PSF of the microlens and then the centroids of each spot were determined using a center of mass algorithm with an iterative window size(33), allowing high-precision, robust estimation even for asymmetric spot patterns. The reliability weight of each spot depends on its signal-to-background ratio ***W*** = Diag(*w*_1_ ⋯ *w_N_*, *w*_1_ ⋯ *w_N_*). The spot’s displacement from the reference pattern ***S***_Sref_ calculated as Δ***S*** = ***S***_all_ – ***S***_Sref_ represents the sample-induced wavefront distortion. Then the additional corrective pattern of the DM can be computed by minimizing the total aberration as follows:

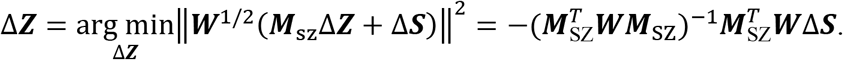

The corrective pattern of the DM for full AO correction is ***Z***_full_ = ***Z***_sys_ + Δ***Z***. Note that all the corrective wavefronts shown in this work represent the corrections for sample-induced aberrations.

### Direct wavefront sensing by line scanning along the microvessels in a 3D subvolume

To measure aberrations at depth, we employed the NIR guide star generated by exciting Evans Blue within the microvasculature in a small 3D volume (~ 60×60×60 μm^3^)). In brief, we performed multiple line scans along the microvessels while integrating the guide star signal on the SHWS. We repeated this procedure at two or more adjacent planes (depth range < 60 μm) and obtained a series of SHWS images (**Fig. 1f-g)**. Because of the large optical scattering induced by the skull and brain tissue, each raw SHWS image displayed distorted focus patterns in various cells of SHWS, yielding errors in aberration correction (**Fig. S6**). To solve this problem, we summed all the SHWS images acquired at different depths and found that the complex irregular and asymmetrical patterns were averaged out in most cells. Further, for those spots with a low signal-background-ratio, we averaged each cell of SHWS with its four nearest neighbors with a weight of 0.25. This wavefront reconstruction algorithm yields a high-quality SHWS image and a single focus stands out in each cell, allowing more accurate determination of the average aberrations in the scattering biological samples (**Fig. S6**).

### Animal preparation

Four transgenic mouse lines: Thy1-YFP (Tg(Thy1-YFP)HJrs/J)(34), Cx3Cr1-GFP (B6.129P2(Cg)-Cx3cr1^tm1Litt^/J)(35), Thy1-YFP/Cx3Cr1-GFP and APP-PS1/Cx3Cr1-GFP were used in this study. Thy1-YFP/Cx3Cr1-GFP mice were generated by crossing Thy1-YFP mice with Cx3Cr1-GFP mice, and APP-PS1/Cx3Cr1-GFP mice were obtained by crossing APP-PS1 (Tg (APPswe, PSEN1dE9)85Dbo) mice with Cx3Cr1-GFP mice. All the animal procedures conducted in this work followed an animal protocol approved by the Animal Ethics Committee of HKUST.

Mice (>6 weeks) were anesthetized by intraperitoneal (i.p.) injection of ketamine/xylazine mixture (10μL/g) before surgery. After the skull was exposed by performing a midline scalp incision, a scalpel was used to remove gently the periosteum attached to the skull. Then a custom-designed rectangular head plate with a circular hole was centered on the right hemisphere and sealed onto the skull by applying a small amount of cyanoacrylate adhesive to the perimeter of the hole. Dental acrylic was then applied to the exposed skull surface to fill the gap between the head plate and skull. After the dental acrylic became dry and hard, the mice were mounted on a head-holding stage with angle adjusters (NARISHIGE, MAG-2) and placed under a stereomicroscope for surgical preparation of either a thinned skull or optical-cleared skull window.

#### Thinned skull window

The thinned skull preparation is slightly modified from a previous protocol(4). Briefly, a 500-μm carbon steel burr attached to a rotatory high-speed drill was used to gently thin a circular region (2.0-2.5mm in diameter) with the center at stereotactic coordinate (3mm, 3mm) laterally and posterior to the bregma point. After removing the majority of the middle spongy bone, a micro surgical blade (no. 6961, Surgistar) was used to carefully thin the skull further to about 40-50 μm. Surface irregularities were reduced by occasionally changing the thinning direction of the surgical blade. Finally, a biocompatible sealant mixture (Kwik-Cast, WPI) which can be peeled off before the imaging experiment was applied to cover the thinned skull window.

#### Optical clearing skull window

The reagents used for optically clearing the skull include: 10% EDTA disodium (D2900000, Sigma-Aldrich), 80% glycerol (G5516, Sigma-Aldrich) and USOCA (consists of S1 and S2)(7–9). S1 is prepared by dissolving urea (Sinopharm, China) in 75% ethanol at a 10:3 volume-mass ratio. S2 is a sodium dodecylbenzenesulfonate (SDBS) prepared by mixing NaOH solution (0.7 M) with dodecylbenzenesulfonic acid (DDBSA, Aladdin) at a 24:5 volume-mass ratio. The skull optical clearing procedure follows the method described in previous reports(7–9). Briefly, the exposed skull was first treated with S1 for about 20 min, with a clean cotton swab gently rubbing the skull surface to accelerate the clearing process. Then the S1 was removed using a cotton ball and replaced with S2 for a further 5 min. After the S2 was removed, 10% EDTA was dropped onto the skull for another 20 min and then replaced with 80% glycerol. Finally, a thin layer of plastic wrap was used to cover the cleared skull to separate the immersion medium (water) from glycerol during *in vivo* imaging.

### *In vivo* imaging

Mice were anesthetized with ketamine/xylazine and received a retro-orbital intravenous injection of Evans Blue (10ug/g; E2129, Sigma-Aldrich) 30 minutes before imaging to label the lumen of blood vessels. The APP-PS1/Cx3Cr1-GFP mice were also i.p. injected with MeO-X04 (5.0 mg/kg, 10% DMSO, 90% PBS) 2 hours before imaging to label amyloid deposits in the brain. Before fluorescence imaging, the skull window was aligned precisely perpendicular to the objective axis by adjusting the angles of the head-holding stage guided by second-harmonic generation imaging of bone collagen.

For two-photon imaging of neurons (YFP), microglia (GFP) and blood vessels (Evans Blue), the femtosecond laser was tuned to 920 nm and the post-objective excitation power ranged from 20-200 mW depending on the imaging depth. To image amyloid plaques in the APP-PS1/Cx3Cr1-GFP mice, another femtosecond laser tuned to 800 nm was used to excite the MeO-X04 fluorescence with an incident power of less than 30mW at the skull surface. For wavefront sensing, the nonlinear fluorescence guide star was created by scanning the 920 nm laser over a small FOV (30×30μm^2^) or selectively choosing a small vessel via multiple line scanning. Detailed imaging and wavefront sensing parameters are listed in **Table S1**.

### Laser-mediated microsurgery

To perform precise and efficient microsurgery using the femtosecond laser, the sample-induced aberration for the ablation site was first measured and compensation applied. For laser dendrotomy, a 920 nm laser with an average power of 400 mW was focused on the dendritic shaft for 1-2s, the actual exposure time being controlled by the feedback signal of newly-created fluorescence during the multiphoton ionization process(36, 37). For single spine ablation, a 920 nm laser with an average power of 200-300 mW was focused on the dendritic spine for 2s.

### Spectra unmixing of GFP and YFP signal

We used Thy1-YFP/Cx3Cr1-GFP mice to study the interaction between neurons and microglia. Because the emission spectra of YFP and GFP are very close, we designed a simple algorithm using a linear model to distinguish the two components. To image GFP and YFP, D3 was replaced with dichroic beam splitter FF518-Di01-25X36 and F2 and F3 were both the band-pass filter FF01-525/50 (**Fig. S1**). Therefore, the detection bands of the two PMT channels are 518-550 nm and 500-518 nm respectively, corresponding to the emission peaks of YFP and GFP. Assuming that the fluorescence brightness of YFP and GFP at the imaging location are *C_Y_* and *C_G_*, and the detected signal intensities of PMT1 and PMT2 are *I*_1_ and *I*_2_, we have the following equations:

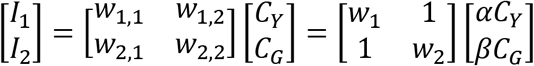

The parameters *w*_1_ and *w*_2_ can be calibrated from the image locations where only YFP labelled neurons or GFP labelled microglia exist. The unmixed normalized signal intensity for YFP and GFP can be represented by *αC_Y_* and *βC_G_*, where:

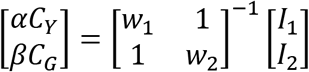

By using this method, we can unmix the fluorescence signal of YFP-labelled neurons and GFP-labelled microglia.

### Image analysis

The images were processed in Matlab (Mathworks) or ImageJ (NIH)(38). Several algorithms for images registration were used to mitigate motion artifacts depending on the level of the animal motion itself. For most imaging conditions when the animal motion was negligible, only a single frame was captured per slice for the stack images. The stack images were registered using the rigid-body transformation provided by the stackreg plugin(39) in ImageJ. When animal motion became apparent and inter-frame artifacts appeared, the imaging speed was increased and several frames were acquired per slice for the whole stack images (**Table S1**). Image registration was performed on sequential frames for each slice using the turboreg plugin(39) in ImageJ to correct the rigid motion artifact, or with the hidden Markov model algorithm(40) to correct within-frame motion artifacts.

For mosaiced images of microglia and blood vessel, multi-tile subimages were captured with predefined positions and then stitched together to form the mosaic image using the Grid/Collection Stitching(41) plugin in ImageJ.

## Acknowledgments

This work was supported by the Hong Kong Research Grants Council through grants 662513, 16103215, 16148816, 16102518, T13-607/12R, T13-706/11-1, T13-605/18W, C6002-17GF, C6001-19E, N_HKUST603/19 and the Innovation and Technology Commission (ITCPD/17-9), and the Area of Excellence Scheme of the University Grants Committee (AoE/M-604/16, AOE/M-09/12) and the Hong Kong University of Science & Technology (HKUST) through grant RPC10EG33.

## Author contributions

C.C., Z.Q., D.Z., N.Y.I. and J.Y.Q. conceived of the research idea. C.C. and Z.Q. designed and conducted the experiments and data analysis. Z.Q., S.H. and C.C. built the AO two-photon imaging system. C.C. carried out the surgery with the assistance of S.L., S.F.L. and W.W.. Finally, Z.Q. and C.C. took the lead in writing the manuscript with inputs from all other authors.

## Supplementary Figures

**Fig. S1.**
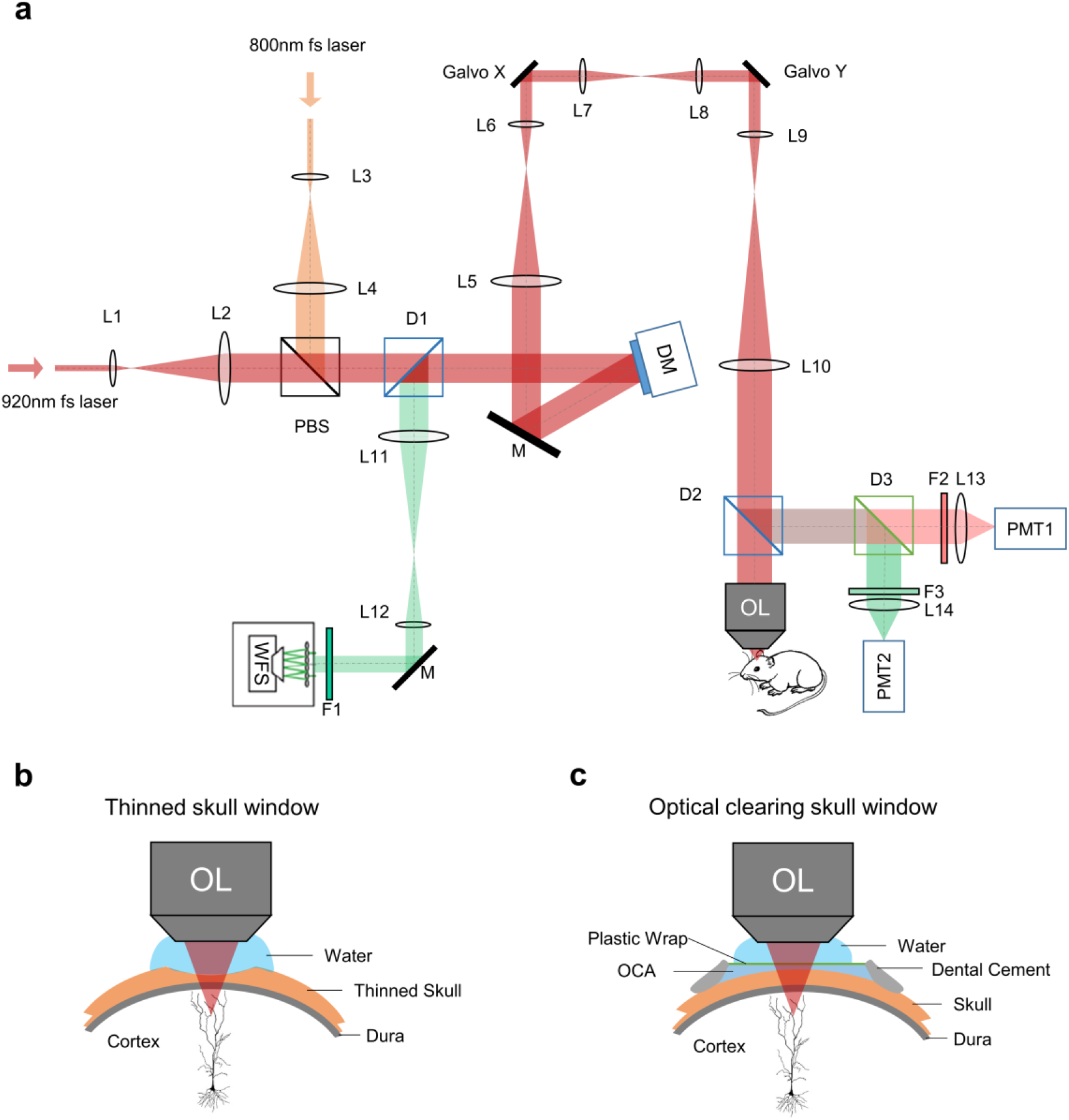
Schematic diagrams of our AO two-photon microscope system and minimally invasive skull windows. (a) Schematic of adaptive optics two-photon microscope setup. L1-L14: lenses; OL: objective lens; D1-D3: dichroic mirrors; F1-F3: filters; M: mirrors; DM: deformable mirror; WFS: wavefront sensor; PMT1-2: photomultiplier tubes. (b) Schematic of thinned skull window. (c) Schematic of optical clearing skull window. OCA: optical clearing agents.

**Fig. S2.**
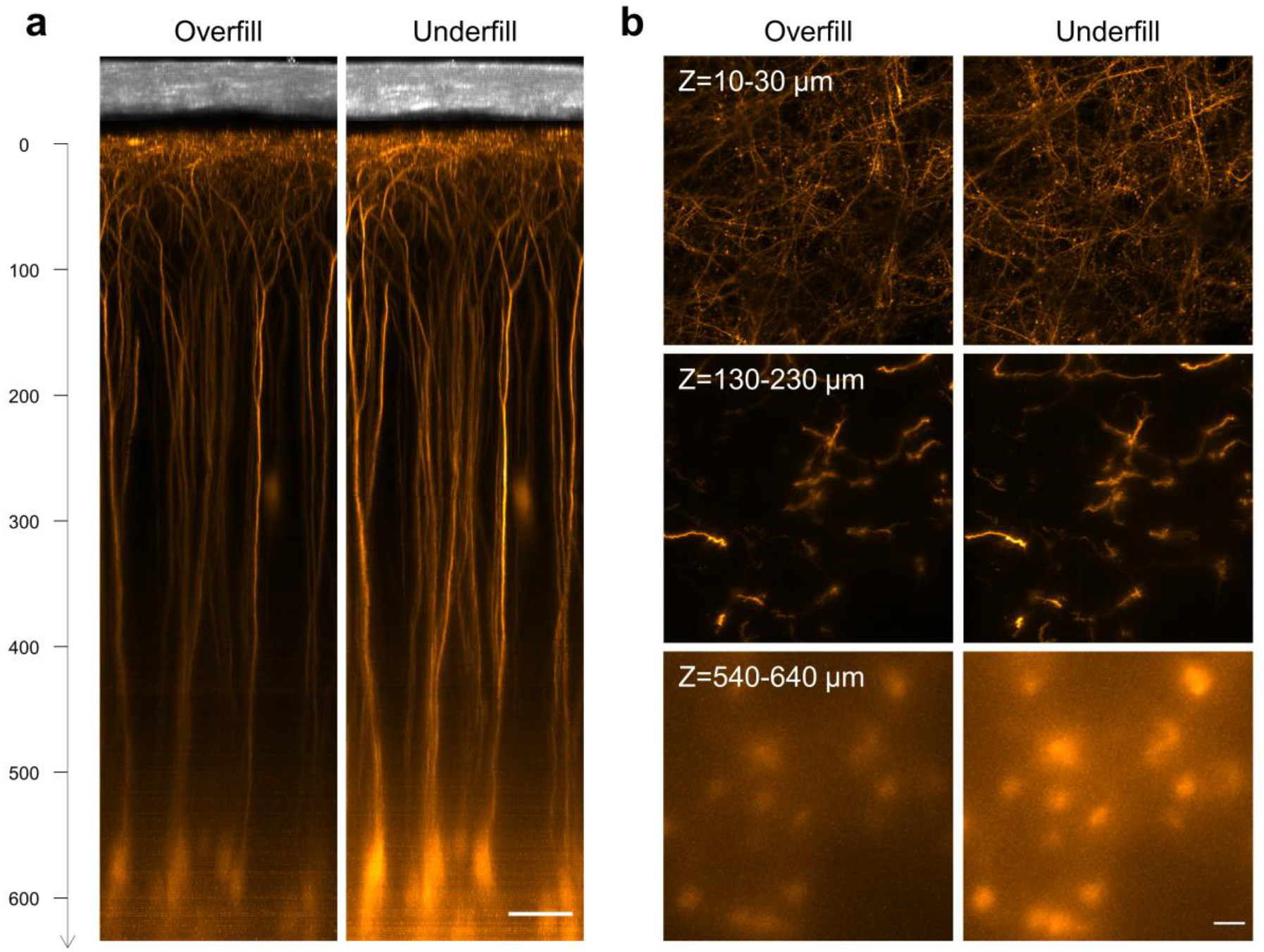
The use of an underfilled objective improved the excitation efficiency for transcranial deep brain imaging. (a) xz maximum-intensity projection (MIP) images of the pyramidal neurons in Thy1-YFP mice through a 50-μm-thickness thinned-skull window acquired using the overfilled (left) and underfilled (right) objective. The imaging conditions including excitation power, pixel size and frame rate were identical at the same depth for both configurations. Scale bar: 50 μm. (b) xy MIP (top row: 10-30μm; middle row: 120-230 μm; bottom row: 540-640 μm) of the images in (a). Scale bar: 20 μm.

**Fig. S3.**
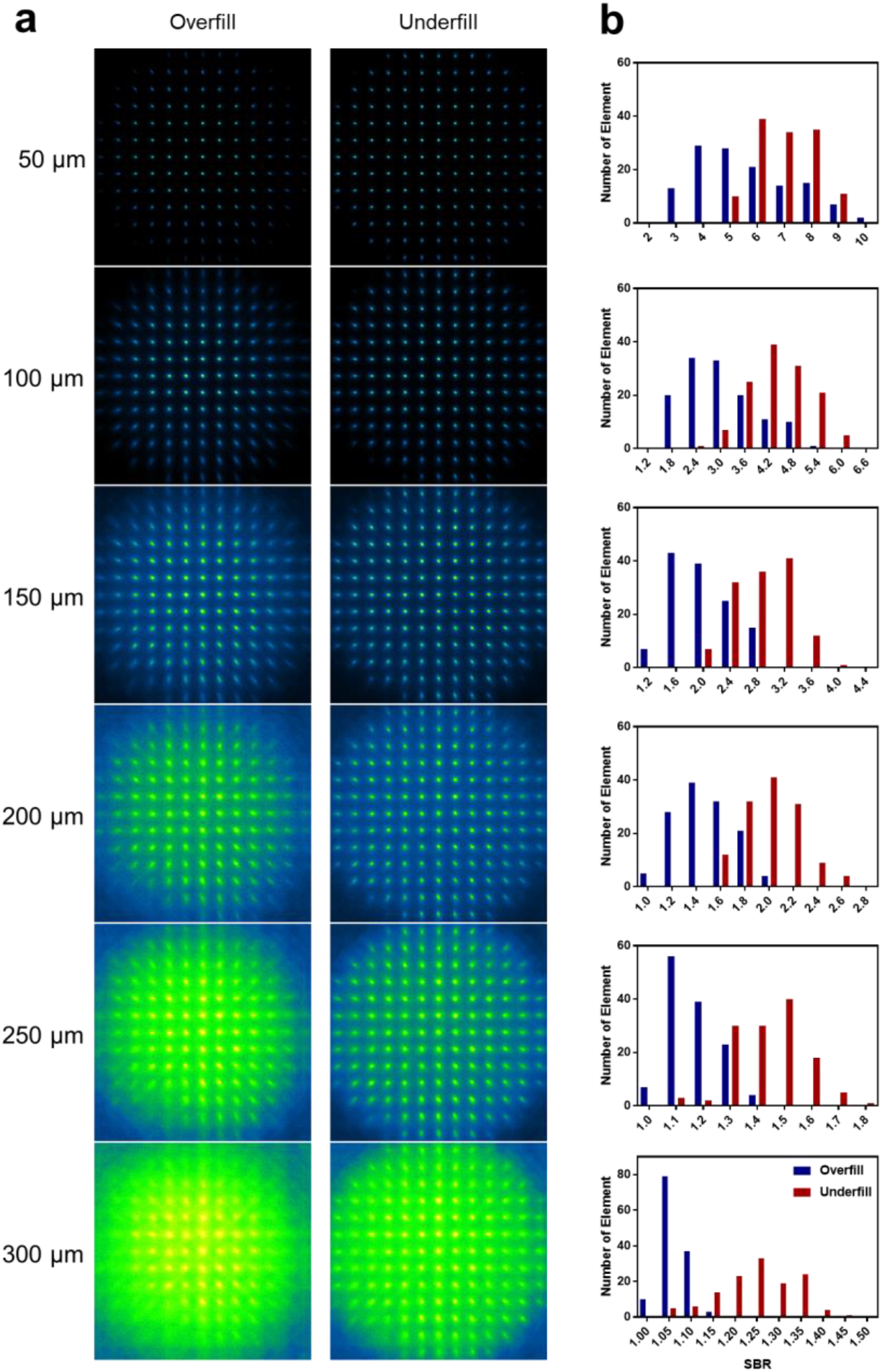
Comparison of guide star images on SHWS using an overfilled or underfilled objective. (a) Guide star images of YFP fluorescence in Thy1-YFP mice when the objective was overfilled (left column) and underfilled (right column) at different depths below the thinned skull. The guide star images were measured at the same location as Fig. S2a. (b) Histograms of the signal-to-background ratio (SBR) of the guide star images at different depths when the objective was underfilled (red) and overfilled (blue). The total element number of the SHWS is 129. The total element number of the SHWS is 129. The SBR was defined by the average intensity of 4×4 pixels around the spots center to that of the remaining pixels in each cell of SHWS. We found that when the SBR was less than 1.2, the fitting of spot center would be inaccurate and thus that spot was defined as a bad spot.

**Fig. S4.**
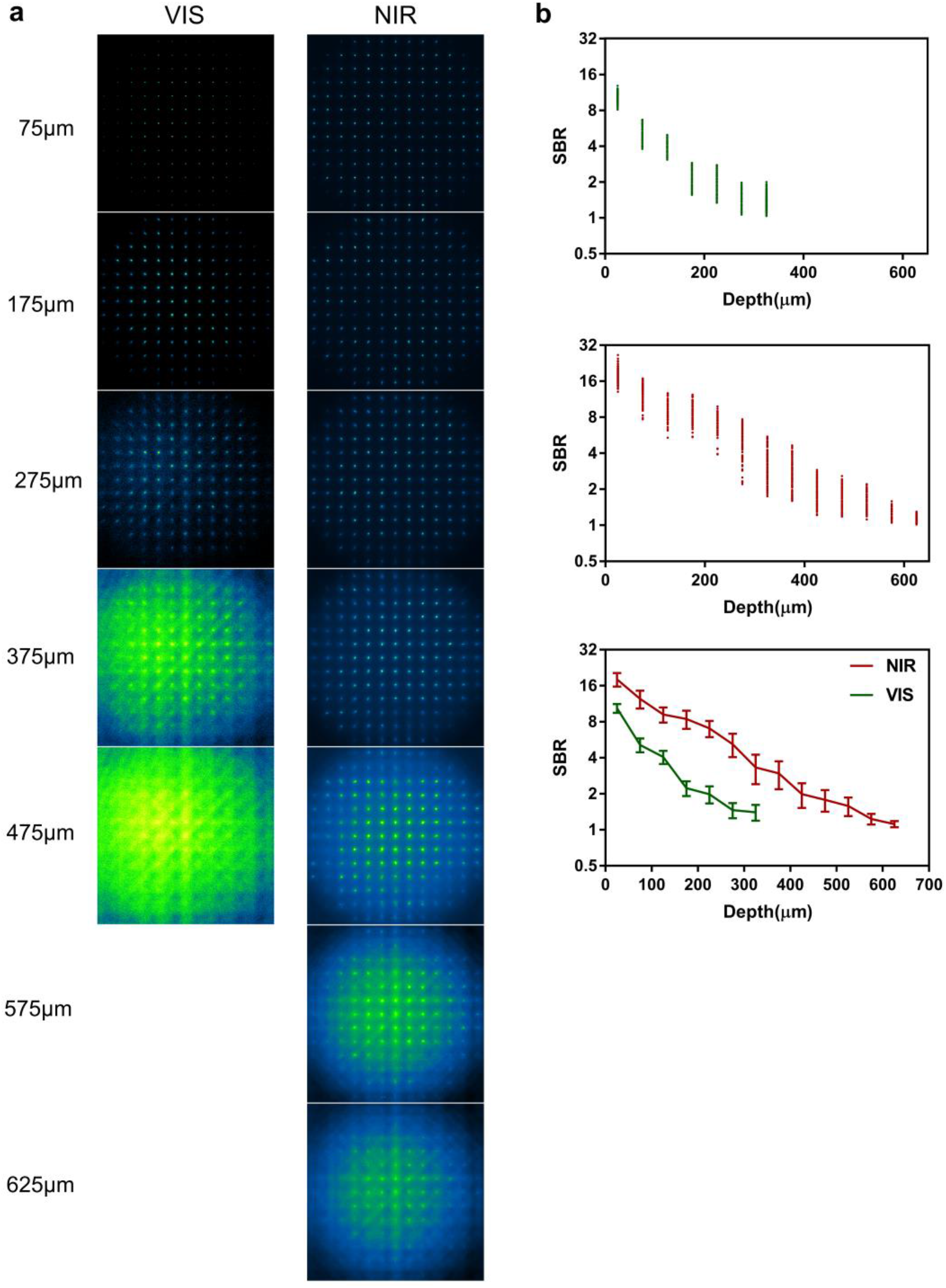
Comparison of the visible and NIR guide star generated by two-photon excitation of YFP and Evans blue in the mouse cortex through a thinned-skull window. (a) Typical guide star images on SHWS at different depths by two-photon excitation of YFP labelled pyramidal neurons (left) and Evans blue labelled microvessels (right) at the same location. (b) The signal-to-background ratio of the visible and NIR guide star.

**Fig. S5.**
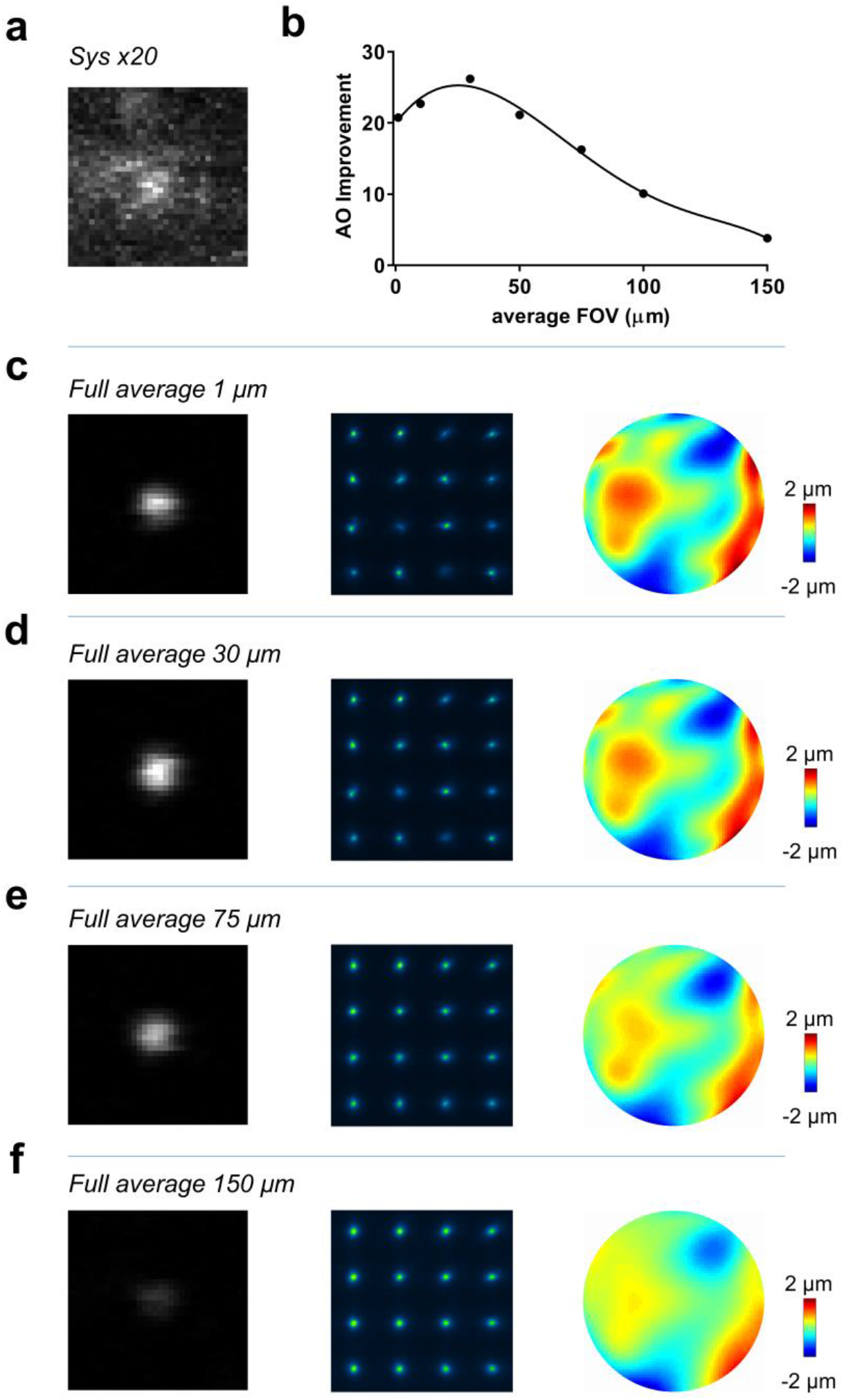
Characterization of the AO corrective FOV through a thinned-skull window. (a) xy MIP image of a 200-nm-diameter bead 400 μm beneath the thinned-skull (50 μm in thickness) with system correction alone. The image brightness was enhanced 20-fold to visualize the details. (b) Improvement of fluorescence intensity of the bead located at the FOV center by AO correction with guide star signals averaged over a square field ranging from 1 μm to 150 μm per side. (c-f) AO corrected xy MIP images of the fluorescent beads (left column), corresponding guide star images (middle column) and corrective wavefront (right column) when the guide star signal is averaged over 1 μm (c), 30 μm (d), 75 μm (e) and 150 μm(f).

**Fig. S6.**
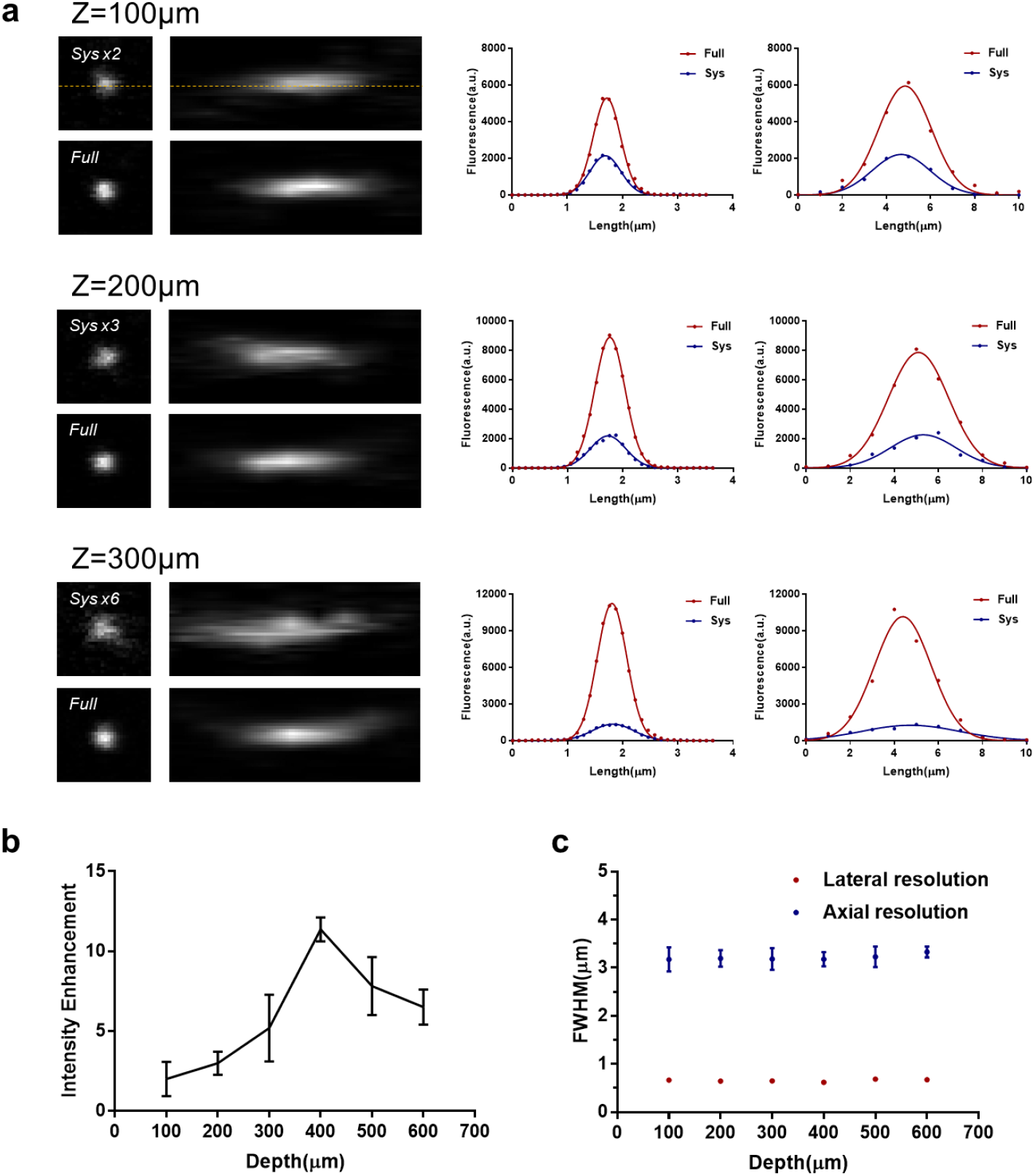
*In vitro* characterization of aberrations of a thinned-skull window. (a) Left column: lateral and axial PSF measured with 200 nm diameter fluorescent beads at different depths with system and full AO correction. The images with system correction were enhanced to visualize details. Middle column: lateral intensity profile along the dashed line with system (blue) and full (red) AO correction. Right column: axial intensity profile with system and full AO correction. Full AO correction was performed by averaging the guide star signal over 30×30 μm^2^. (b) The improvement of fluorescence intensity by full AO correction at different depths. (c) The lateral (red) and axial (blue) resolution after full AO correction.

**Fig. S7.**
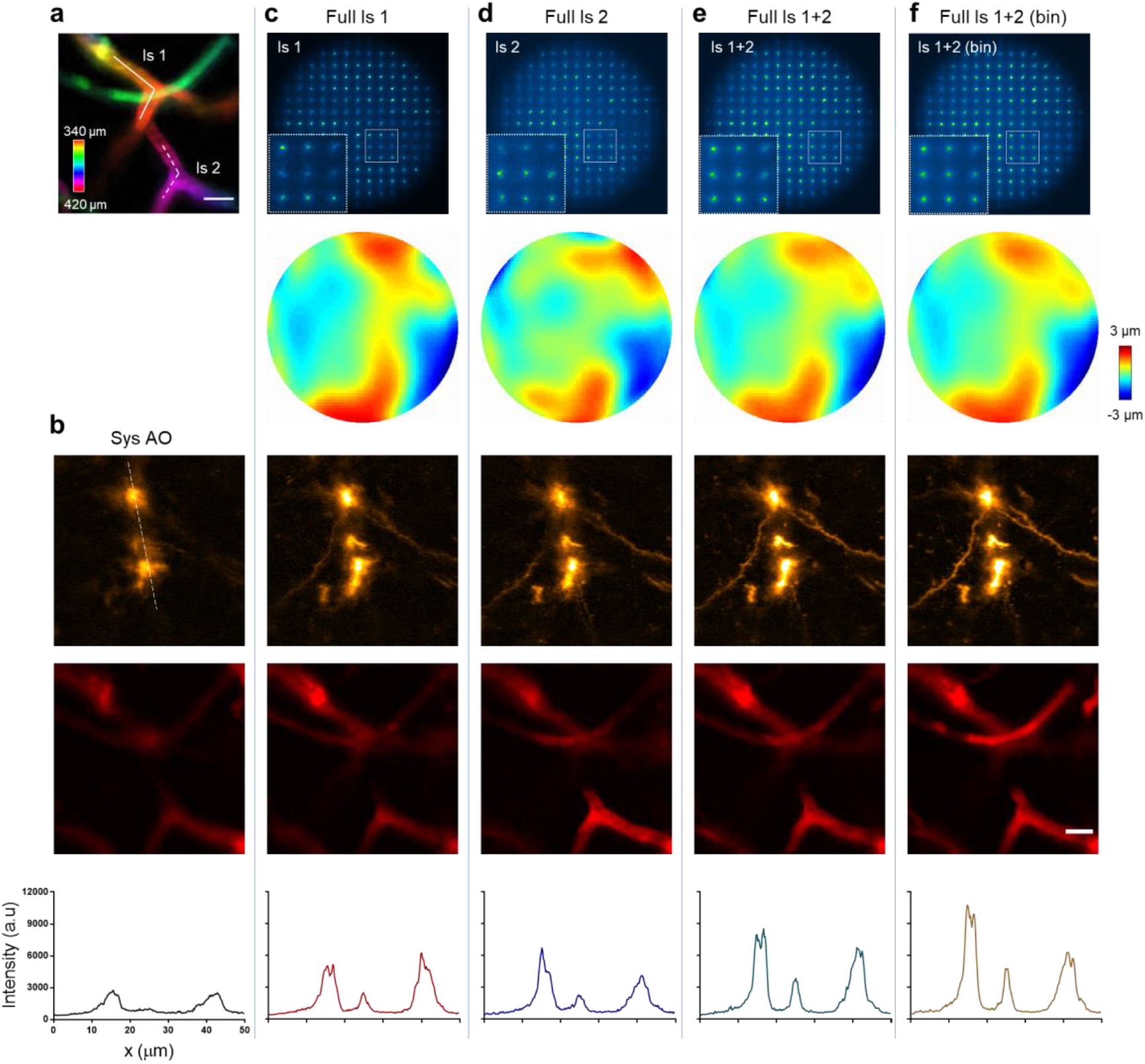
Direct wavefront sensing algorithm for through-skull imaging of the brain. (a) 3D distribution of the vasculature for direct wavefront sensing using a NIR guide star. Two segments of vessels (solid line and dashed line labelled with ls 1 and ls 2) at different depths were line scanned for wavefront measurement. (b) xy MIP images (Z = 340-420 μm) of neuron (top) and microvasculature (middle) and intensity profile along the dashed line (bottom) with only system aberrations corrected. (c-f) 1^st^ row: guide star images on the SHWS, the inset shows a magnified view of the box region; 2^nd^ row: the corresponding corrective wavefront on the DM; 3^rd^ row: MIP images of neurons after correction; 4^th^ row: MIP images of microvessels after correction; 5^th^ row: intensity profile along the dashed line in (top panel of (b)). The AO corrections in (c-f) were based on direct wavefront measurement with only ls 1 (c) or ls 2 (d), the sum of ls 1 and ls 2 (e) and our algorithm by summing ls 1 and ls 2 and subsequent filtering (f).

**Fig. S8.**
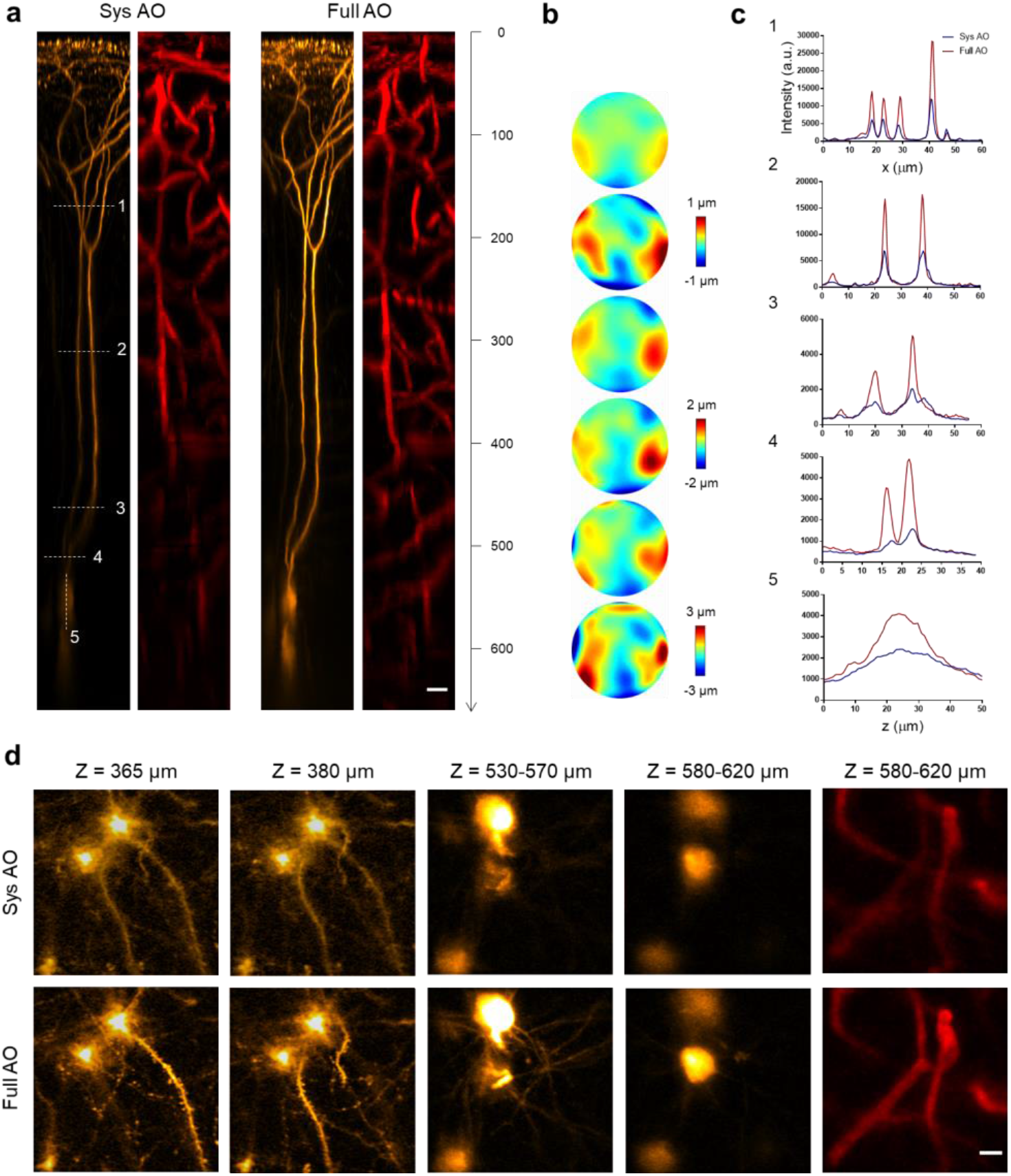
AO recovers high-resolution imaging of the cortex through a thinned-skull window. (a) xz MIP images of the pyramidal neurons (orange) and microvasculature (red) in Thy1-YFP mice through a thinned-skull window (50 μm in thickness) with system correction only (left) and full AO correction (right). AO correction was performed every 50 μm deep. Scale bar: 20 μm. (b) Representative corrected wavefront of the DM at depths (Z = 100, 200, 300, 400, 500 and 600 μm) used in (a). (c) The intensity profile of the dashed line in (a) with system (blue) and full (red) AO correction. (d) xy MIP of the stack images in (a). Orange: neuron; red: microvasculature. Scale bar: 10 μm.

**Fig. S9.**
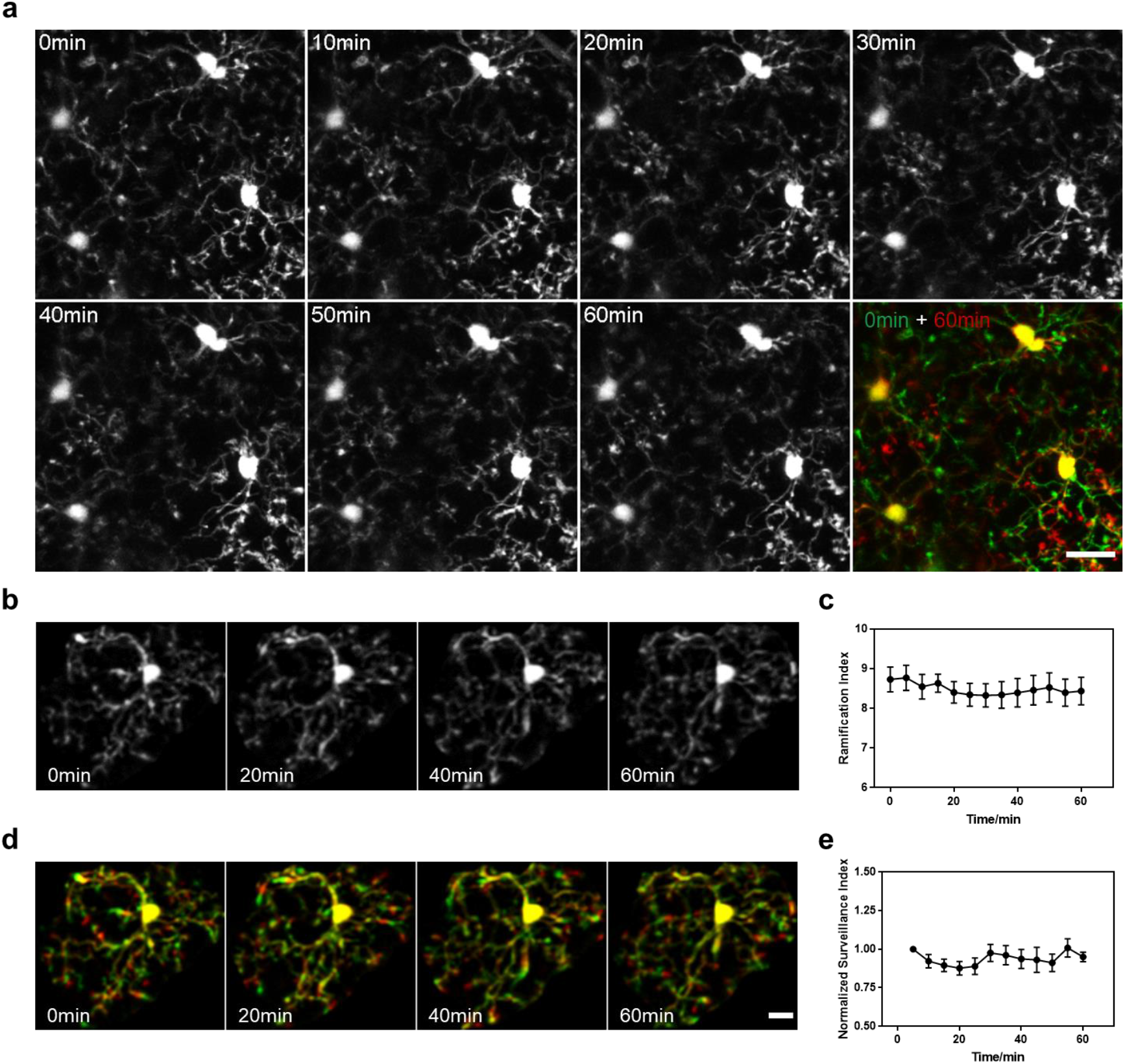
Investigation of microglial inflammation of a thinned-skull window. (a) Time-lapse images showing that microglia remain stable after preparing the thinned-skull. Scale bar: 20 μm. (b) Representative magnified images showing that microglia remain highly ramified under the thinned-skull window. (c) Changes in the microglial ramification index at different times. (d) Merged images (green and red) of two consecutive time points at 5 min interval, showing microglial process movement during surveillance (green: retracted, red: extended) at different times. Scale bar: 10 μm. (e) Changes of microglial surveillance index at different times. Data were normalized to initial time point.

**Fig. S10.**
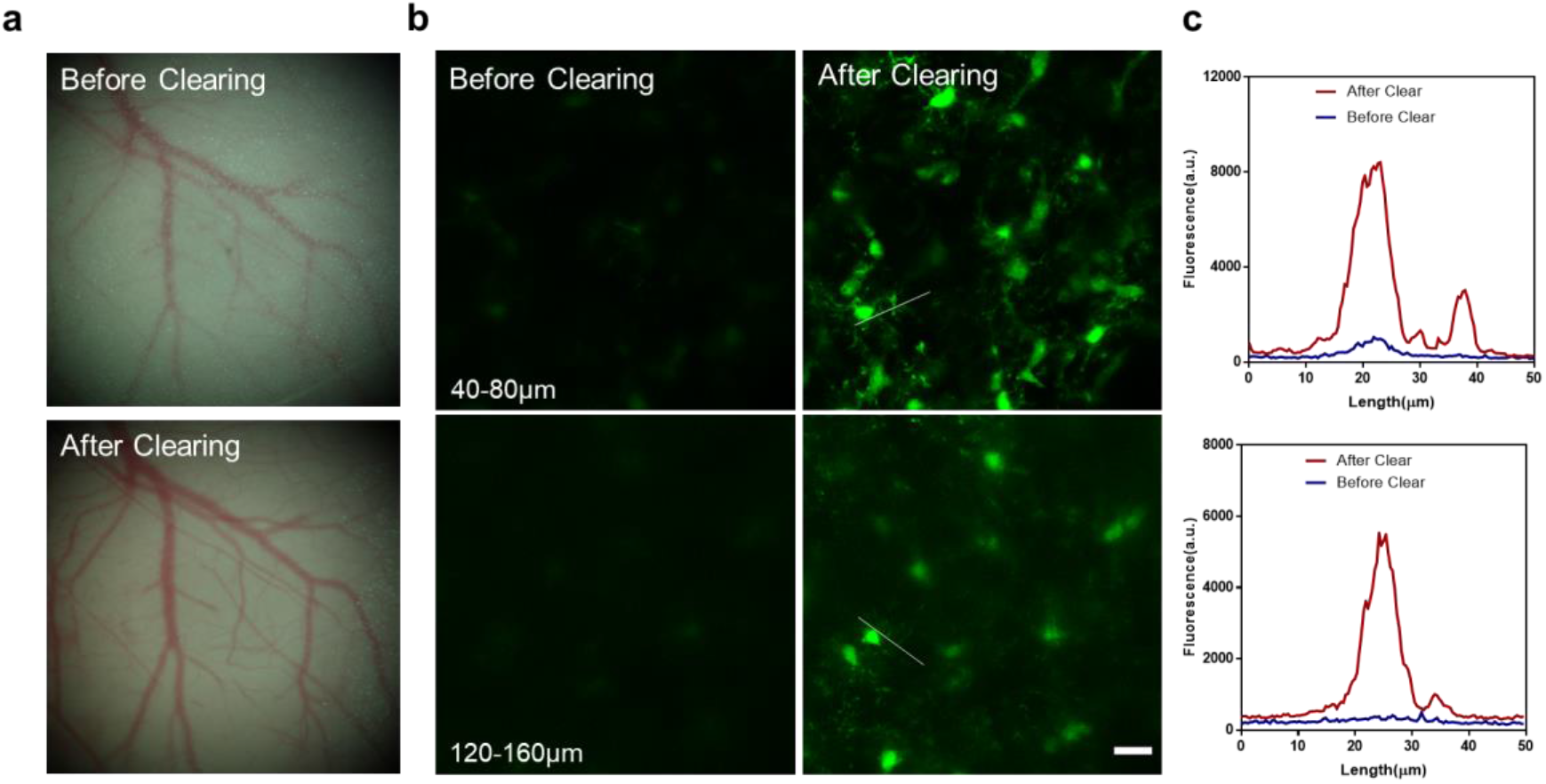
*In vivo* imaging of the brain through an optical clearing window. (a) Bright-field image before (top) and after (bottom) optical clearing. (b) xy MIP of a two-photon image of the GFP labelled microglial in the Cx3Cr1-GFP mice before (left column) and after (right column) optical clearing. Imaging depths: top row: Z = 40-80 μm; bottom row: Z = 120-160 μm. (c) Intensity profile along the lines in (b).

**Fig. S11.**
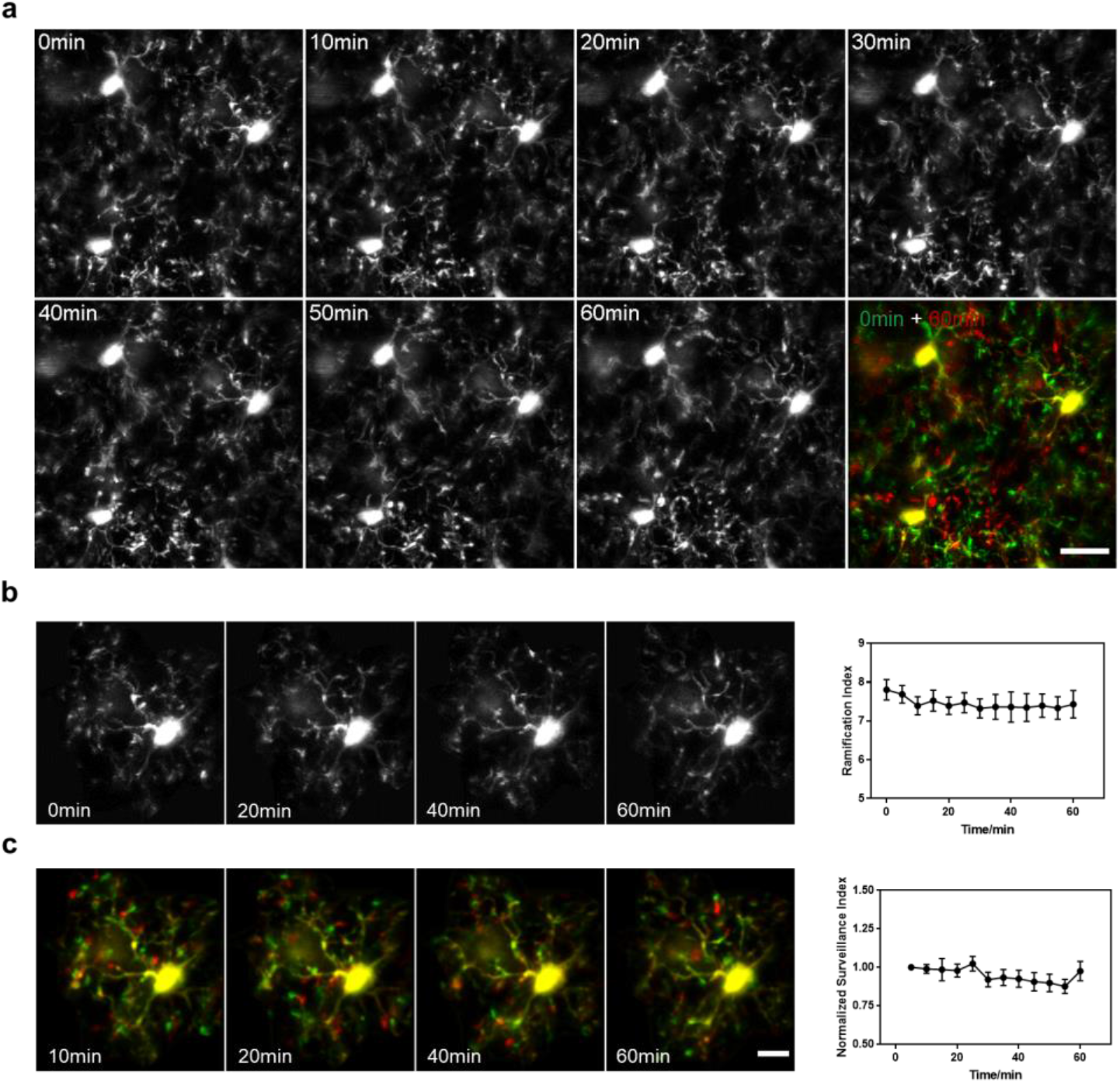
Investigation of microglial inflammation of an optical clearing window. (a) Time-lapse images showing that microglia remain stable after optical clearing of the skull. Scale bar: 20 μm. (b) Representative magnified images showing that microglia remain highly ramified under the thinned-skull window. (c) Changes in the microglial ramification index at different times. (d) Merged images (green and red) of two consecutive time points at 5 min interval, showing microglial process movement during surveillance (green: retracted, red: extended) at different times. Scale bar: 10 μm. (e) Changes of microglial surveillance index at different times. Data were normalized to initial time point.

**Fig. S12.**
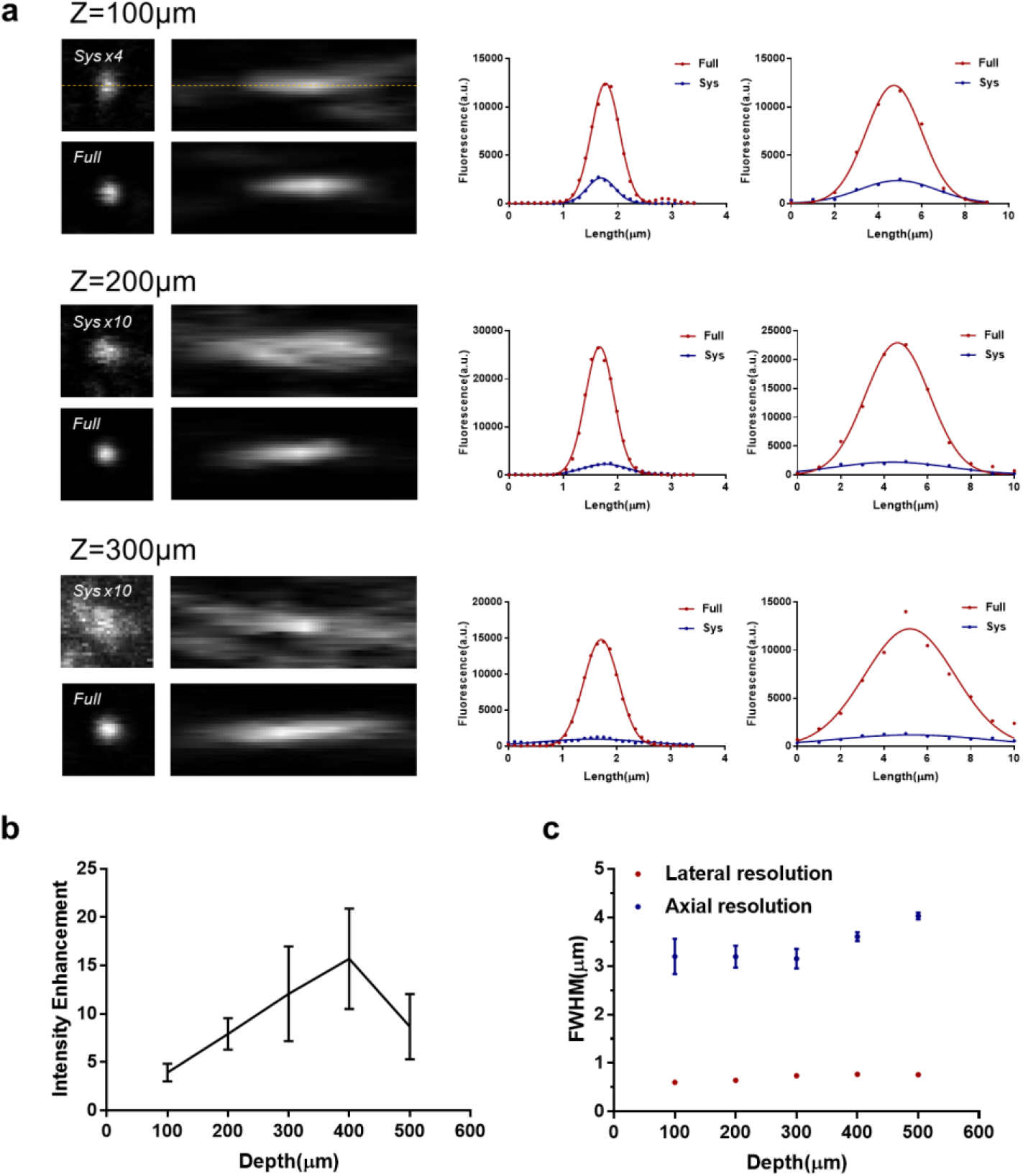
*In vitro* characterization of aberrations of an optical clearing window. (a) Left column: lateral and axial PSF measured with 200-nm-diameter fluorescent beads at different depths with system and full AO correction. The images with system correction were enhanced to visualize details. Middle column: lateral intensity profile along the dashed line with system (blue) and full (red) AO correction. Right column: axial intensity profile with system and full AO correction. Full AO correction was performed by averaging the guide star signal over 30×30 μm^2^. (b) The improvement of fluorescence intensity by full AO correction at different depths. (c) The lateral (red) and axial (blue) resolution after full AO correction.

**Fig. S13.**
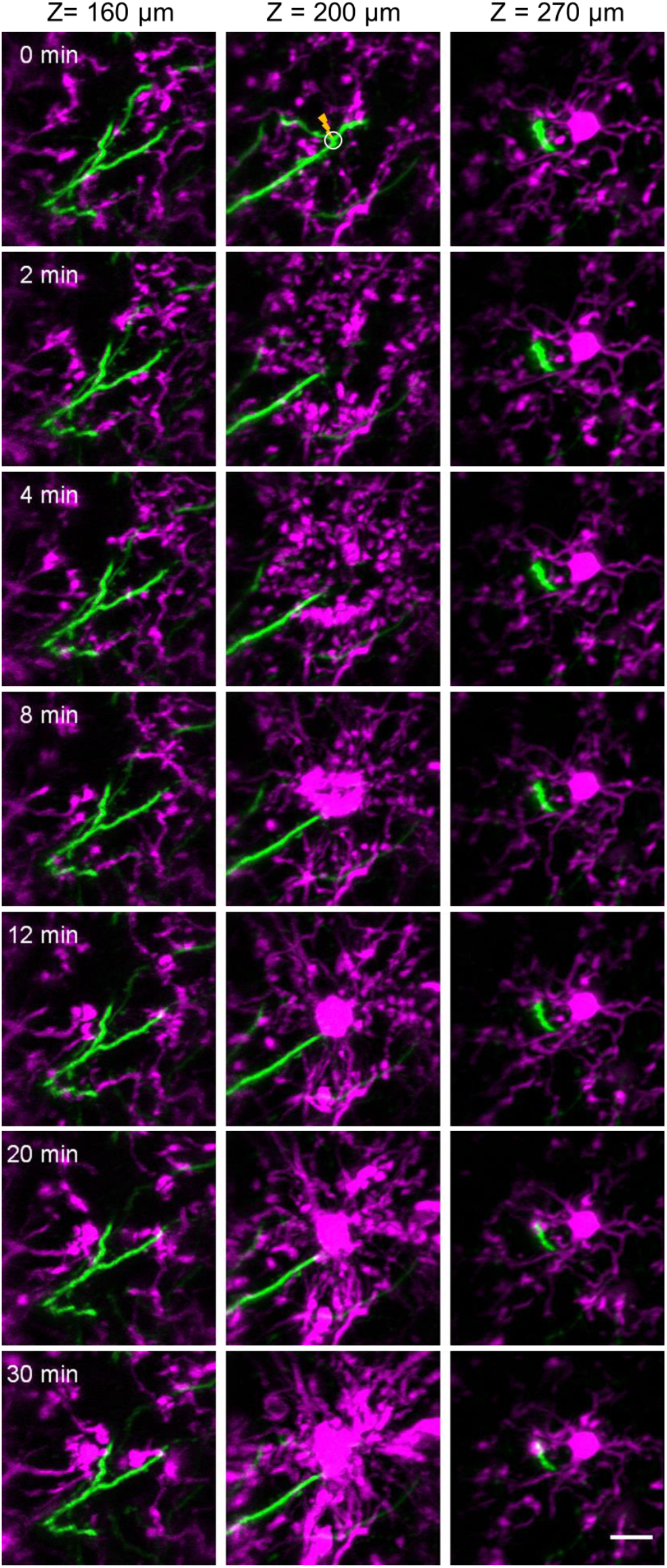
Time-lapse imaging of neurons (green) and microglia (magenta) in response to laser-mediated lesion of the bifurcation of a primary apical dendrite. The solid circle in the top panel indicates the site of the laser injury. Scale bar: 10 μm.

**Fig. S14.**
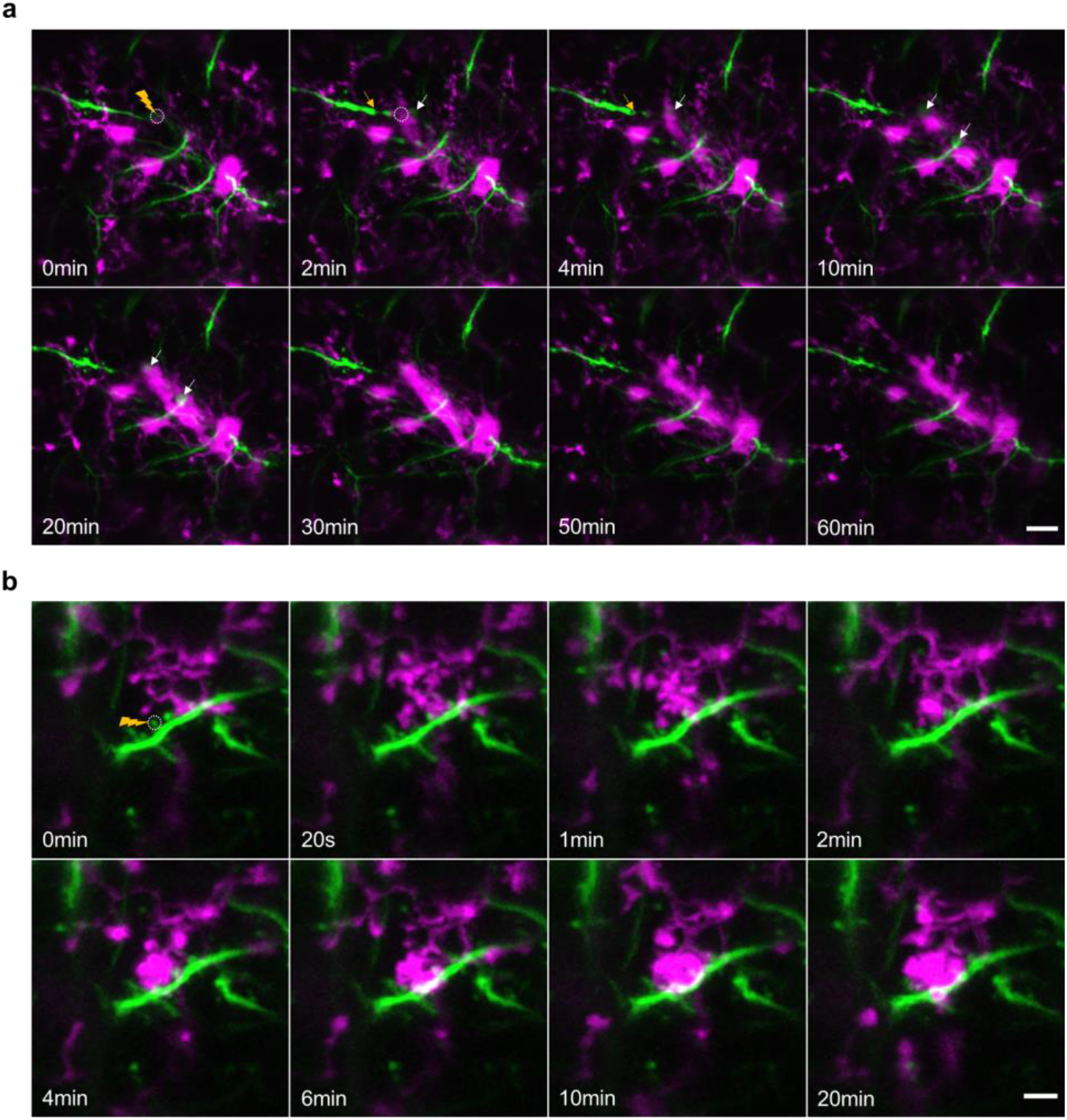
AO enables *in vivo* study of neuron-microglia interaction after laser ablation of the dendritic branch and a single spine. (a) Time-lapse imaging of neuron (green) and microglia (magenta) in response to laser dendrotomy on a tuft dendrite of the layer 5 pyramidal neuron. The injured region is indicated by the dashed circle. Two white arrows indicate the retraction bulb formation at the proximal end of the injured branch, and the yellow arrow indicates the dendritic segmentation at its distal end. The results show that while the injured dendrite undergoes Wallerian-like degeneration at both distal and proximate ends, the nearby microglial process only extended selectively to surround the proximate end of the retraction bulb rather than the distal end. This phenomena may be potentially explained by different signaling mechanisms for distal and proximate dendrites during neuronal degeneration. Scale bar: 10 μm. (b) Time-lapse imaging of the dynamics of microglial processes in response to precise micro-lesion of a single spine (dotted circle in the first frame) without damaging the dendritic shaft and nearby spines. In contrast to laser cutting of the dendrite, this micro-lesion only triggers the activation of a few nearby microglia, whose processes rapidly converged on the ablated spine as soon as 2 minutes after injury. Scale bar: 5 μm.

**Table S1.**
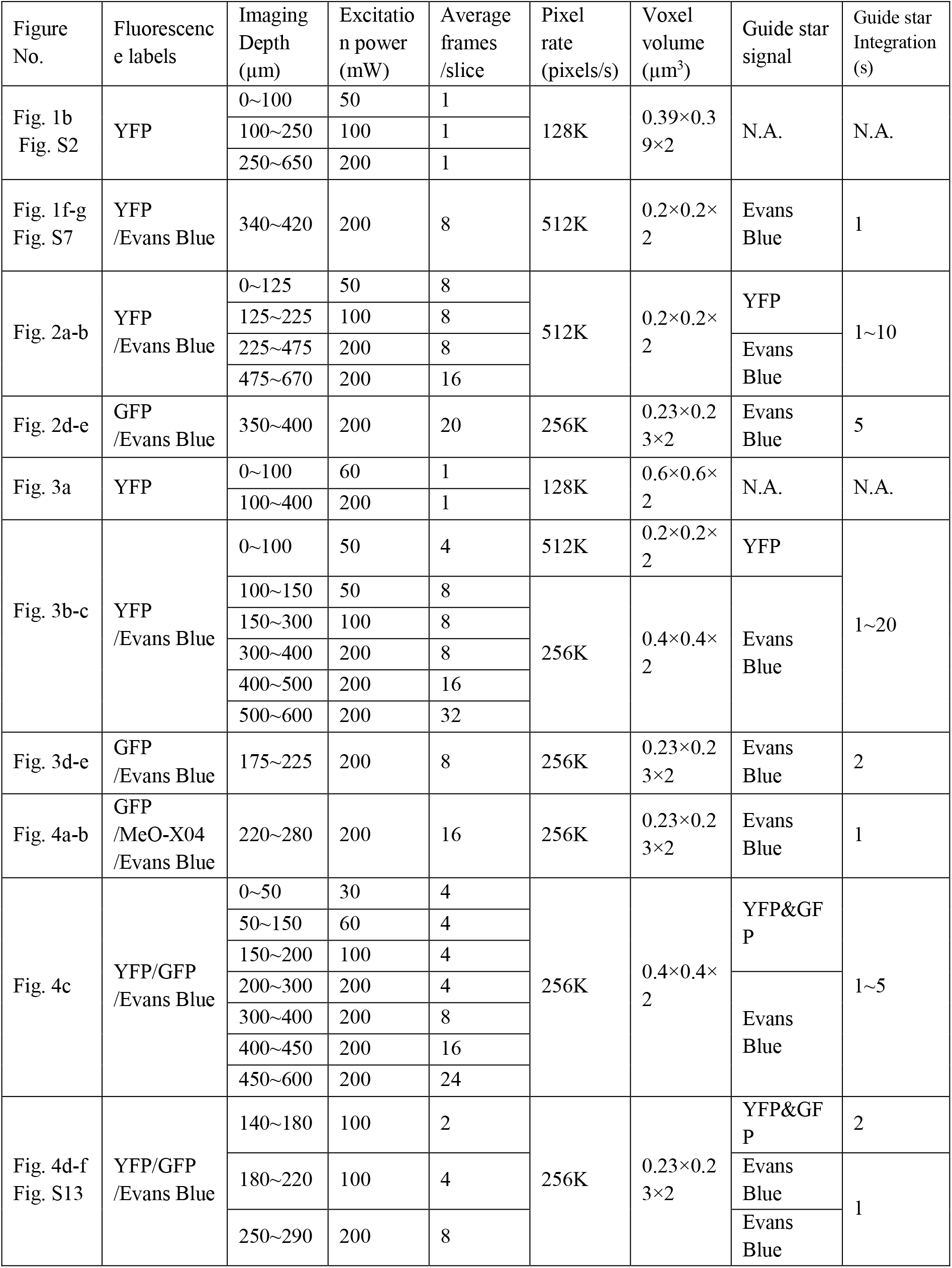
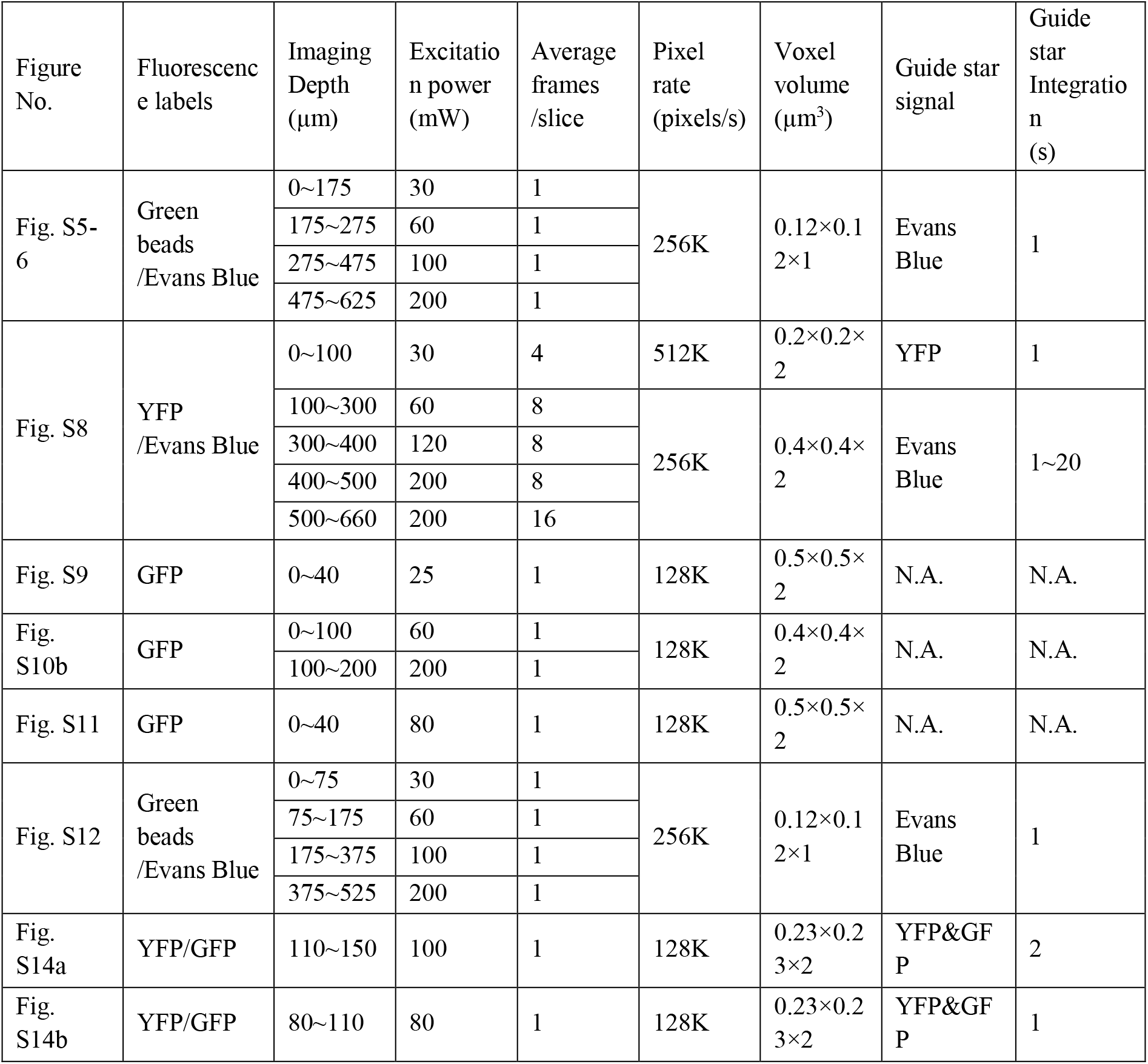
Wavefront sensing and imaging parameters.

**Movie S1 (separate file).** Time-lapse imaging of neuron-microglial interactions following precise laser micro-lesion of bifurcation point of the primary apical dendrite of a layer 5 pyramidal neuron.

**Movie S2 (separate file).** Time-lapse imaging of microglial response to high-precision laser ablation of a single spine.

